# Sex-dimorphic neuroprotective effect of CD163 in an α-synuclein mouse model of Parkinson’s disease

**DOI:** 10.1101/2022.10.17.512526

**Authors:** Sara A. Ferreira, Conghui Li, Ida H. Klæstrup, Zagorka Vitic, Rikke K. Rasmussen, Asger Kirkegaard, Gitte U. Toft, Cristine Betzer, Pia Svendsen, Poul H. Jensen, Yonglun Luo, Anders Etzerodt, Søren K. Moestrup, Marina Romero-Ramos

## Abstract

Alpha-synuclein (α-syn) aggregation and immune activation represent hallmark pathological events in Parkinson’s disease (PD). The PD-associated immune response encompasses both brain and peripheral immune cells, although little is known about the immune proteins relevant for such response. We propose that the upregulation of CD163 observed in blood monocytes and in the responsive microglia in the PD patients is a protective mechanism in the disease. To investigate this, we used the PD model based on intrastriatal injections of murine α-syn pre-formed fibrils (PFF) in CD163 knockout (KO) mice and wild-type littermates. CD163KO females revealed an impaired and differential early immune response to α-syn pathology as revealed by immunohistochemical and transcriptomic analysis. After 6 months, CD163KO females showed an exacerbated immune response and α-syn pathology, which ultimately led to dopaminergic neurodegeneration of greater magnitude. These findings support a novel, sex-dimorphic neuroprotective role for CD163 during α-syn-induced neurodegeneration.

## INTRODUCTION

Parkinson’s disease (PD) is a neurodegenerative disease classically characterized by the loss of dopaminergic neurons in the substantia nigra (SN) and the intraneuronal alpha-synuclein (α-syn) aggregation in Lewy bodies^1^. An immune response has been suggested to parallel, or even precede neurodegeneration in PD, influencing neuronal health ^2^. Microgliosis and a marked pro-inflammatory profile are found in *postmortem* PD brains ^3,4^ and in α-syn-based models ^5–7^. This immune response involves not only brain-resident glia (microglia and astrocytes), but also entails peripheral innate and adaptive immune cells^2^. Interestingly, growing evidence support sex differences in the immune changes associated with PD that might have a significant relevance in the PD risk, presentation and progression ^8–11^. However, little is known about what proteins might be involved in the differential immune response and/or which ones might exert a neuroprotective effect.

The scavenger receptor CD163 is considered a marker for anti-inflammatory macrophages. CD163 expression increases upon anti-inflammatory cytokine IL-10 or glucocorticoid exposure ^12^, but also under certain inflammatory conditions, putatively as a protective mechanism ^13^. Conversely, activation of pro-inflammatory receptors results in CD163 cleavage from the monocytes-membrane (producing soluble (s)CD163), thus associating membrane-bound CD163 loss with pro-inflammatory phenotypes^14^. CD163 uptakes the hemoglobin/haptoglobin complexes for lysosomal degradation^13^. CD163’s specific role in the immune system is still undefined, although studies of CD163 knock out (CD163KO) mice support an anti-inflammatory function ^15,16^. Increased CD163+ cell numbers have been found in PD and Alzheimer’s disease (AD) *postmortem* brains ^17,18^ and in the brain of PD models^19,20^. CD163 expression is lost during microglia development, hence absent in adult surveilling microglia ^21–23^. Therefore, CD163 expression in the brain seems associated with infiltration or otherwise ectopic CD163 upregulation in microglia. Interestingly, recent single cell (sc)RNA sequencing studies show that CD163 upregulation is associated with microglia response during AD and PD in humans ^24,25^.

Our prior data supports a role for CD163-expressing cells in patients with PD and a sex-dimorphic behavior of the CD163 receptor ^26^. Moreover, we have reported that higher percentage of CD163 cells in the blood was associated with lower inflammation in the brain and better putaminal dopaminergic neurotransmission in REM sleep behavior disorder (RBD) patients, suggesting a protective function for the CD163 cells in prodromal stages of PD ^27^. Lastly, we recently showed increased number of CD163 cells and expression levels in the blood of PD patients ^11^. We speculate that the observed increase in CD163 expression is a part of a neuroprotective compensatory mechanism exerted by myeloid cells.

To investigate the significance of the CD163 receptor in the PD-like neurodegeneration we analyzed α-syn neurotoxicity, pathology, and immune alterations in CD163KO mice by injecting recombinant α-syn pre-formed fibrils (PFF) into the striatum of these animals vs. wild type (WT). Additionally, we performed SMARTseq2 on microglia and macrophage populations in the brain to assess the transcriptomic alterations in these immune populations upon CD163 deletion. We hypothesize that CD163 deficiency leads to impaired immune responses to α-syn PFF in a possible sex-dependent manner.

## MATERIALS AND METHODS

CD163KO (CD163^tm1.1(KOMP)Vlcg^) mice and WT (C57BL/6N) littermates (3-4 months old) ^28^ received unilateral intrastriatal injection of murine α-syn pre-formed fibrils (PFF), monomeric α-syn (MONO) or PBS as a control. For histological analysis, mice were sacrificed 1- and 6-months post-surgery (**Supplem. Fig. 1**). Motor behavior was assessed one week before each end-point. For transcriptomic studies, mice received bilateral intrastriatal injection of α-syn MONO or PFF and were euthanized 2 months post-surgery, their brains were dissected, and microglia & macrophage populations were FACS sorted, RNA was purified for sequencing (**Supplem. Fig. 2**). For the *in vitro* experiment, bone-marrow derived macrophages (BMDMs) were isolated from CD163KO and WT mice. All animal experiments were approved and performed under humane conditions in accordance with the ethical guidelines established by the Danish Animal Inspectorate and following EU legislation. See supplementary information for detailed methodological descriptions.

## RESULTS

### Intrastriatal α-syn PFF induced long-lasting motor impairment in CD163KO males

To induce α-syn pathology, murine α-syn PFF (10 µg, average size 33.87nm), α-syn MONO, or control PBS were unilaterally injected into the striatum. Based on our prior observations regarding CD163 in humans with PD, we analyzed sexes separately ^26^. To evaluate whether the genetic deletion of CD163 would lead to differential sensorimotor behavioral changes after α-syn PFF injection, mice were assessed with the challenging beam at 1- and 6-months’ post-injection (p.i.) (**Fig. 1a-i and Supplem. Fig. 3**). At 1-month, we observed a significant increase in the number of steps, errors, errors/step on frame 4 and total errors (the narrowest and most challenging frame), but no changes in time, in all α-syn-injected animals (males and females) when compared to their respective PBS-controls (**Fig. 1a-f, Supplem. Fig. 3a-b**). This suggests that, not only fibrillar α-syn, but also an excess of monomeric protein, is sufficient to induce motor deficits in mice. PFF-CD163KO males had a significantly higher number of errors and errors/step in frame 4 (vs. WT, PBS- and MONO-CD163KO males) (**Fig. 1a-b**) and more total errors (frames 2-4) (vs. PFF-WT and PBS-CD163KO males) (**Fig. 1c**). No changes in errors were found between different genotypes in the female groups (**Fig. 1d-f**). When comparing sexes, PFF-CD163KO males showed significantly higher number of errors in frame 4 than PFF-CD163KO females (**Table 1**, **Fig. 1h**), indicating a sex-specific role for CD163 in the early response to α-syn PFF. Analyzing the temporal evolution in the motor performance most of the readouts at 6 months remained similar to those at 1 month. PFF-CD163KO males still showed a greater number of total errors and total errors/ step than PFF-WT males after 6 months, though no differences between PFF and MONO-males were seen (**Supplem. Fig. 3e**). At 6 months all PFF-injected females took longer to transverse frame 4 than at 1 month (Two-way ANOVA genotype & time, p<0.05), but no differences were seen among the female groups (**Supplem. Fig. 3b, d, f**).

**Fig. 1.**
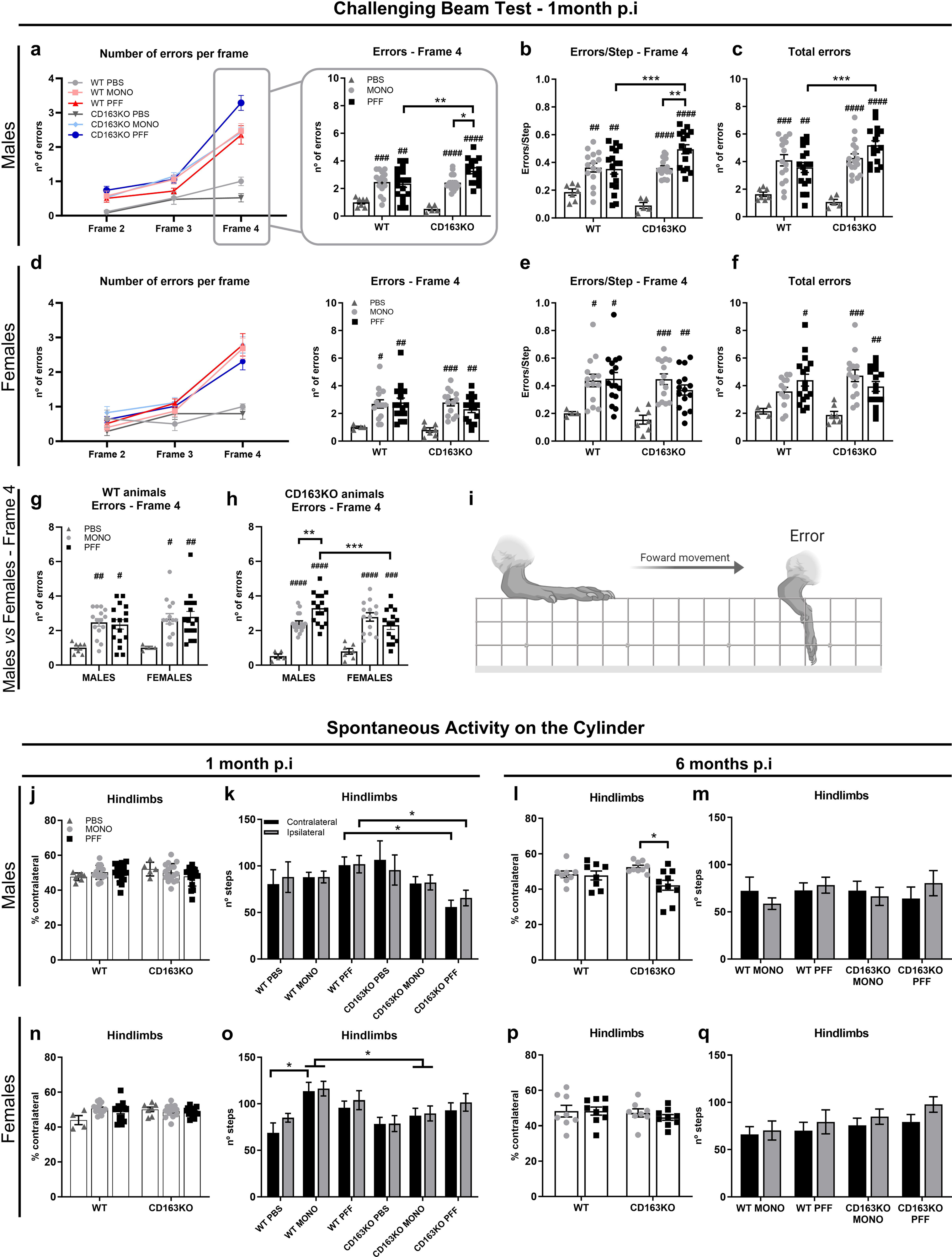
Assessment of motor performance on the Challenging Beam and the Cylinder tests. Top: Line with average and bar graphs with individual points show number of errors per frame, number of errors and errors/step in Frame 4, and total number of errors (sum of frames 2-4,) in the Challenging Beam test, 1-month post-PBS and α-syn MONO/PFF unilateral injection in the Striatum of **a-c)** males and **d-f)** females. **g-h)** Show error differences between males and females in WT and CD163KO animals. **i)** Illustration representing the portrayal of an error during a forward movement (image created with BioRender.com). **Bottom:** Bar graphs with/without individual points represent the contralateral hindlimb use (as percentage of total hindlimbs) and the number of hindlimb steps in the cylinder in **j-m)** males and **n-q)** females. Values are Mean±SEM (n=4-7 (PBS group), 14-17 (MONO/PFF group) for behavior at 1-month; n=8-10 for behavior at 6-months). Statistics: Two-way ANOVA followed by Sidak’s multiple comparison test. P values are given according to the number of symbols e.g. (*p<0.05, **<0.01, ***<0.001, ****<0.0001). # different to corresponding PBS control.

**Table 1.**
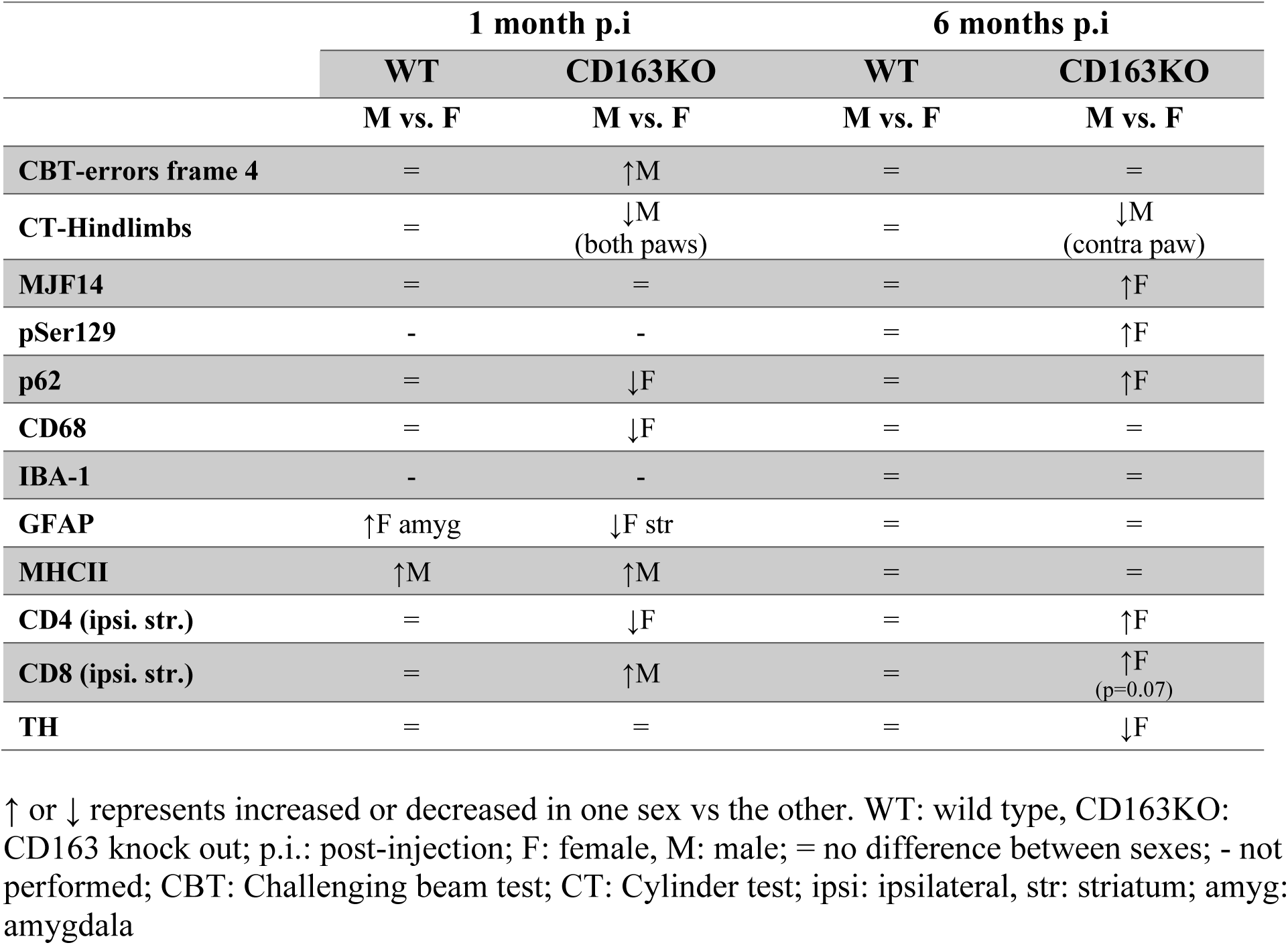
Sex differences in the PFF-injected groups within genotypes.

Evaluation of the spontaneous activity in the cylinder (**Fig. 1j-q**) at 1-month showed a general decrease in hindlimbs use in PFF-CD163KO males (vs. PFF-WT males) (**Fig. 1k).** MONO-WT females had more hindlimbs steps (vs. PBS-WT and MONO-CD163KO females) suggesting certain motoric consequences after α-syn-MONO injection that was affected by the CD163 absence in females (**Fig. 1o)**. When comparing sexes, MONO-WT females and PFF-CD163KO females had higher number of hindlimbs steps (contralateral & ipsilateral) than MONO-WT males and PFF-CD163KO males, respectively (**Table 1, Supplem. Fig. 4a-d**). At 6-months, the total paw use was similar across groups, however, PFF-CD163KO males showed paw-asymmetry by using the contralateral hindlimb significantly less than MONO-CD163KO (**Fig. 1l**). In conclusion, both MONO and PFF α-syn injections lead to motor alterations at short term that appeared distinct and sex-dependent in CD163-deficient mice, but not in WT mice. Collectively, these observations suggest a sex dimorphism in the CD163 system response to α-syn.

### CD163KO females display enhanced α-syn pathology

Accumulation of pathological α-syn aggregates was analyzed using two antibodies: the MJF14, which preferentially binds aggregated α-syn ^29–31^; and an anti-phosphorylated α-syn at Ser129 (pSer129), which is enriched in PD brains ^32^ (**Fig. 2a, Supplem. Fig. 5**). At 1-month p.i, MJF14+ α-syn pathology was present in all PFF-injected groups in the ipsilateral hemisphere of all regions evaluated (striatum, piriform cortex, amygdala, thalamus and SN) and was mostly prevalent in neuropil-like structures (**Supplem. Fig. 5a-d**). No major MJF14+ pathology was seen in the contralateral hemisphere, except few structures in cortex and amygdala (not-shown). After 6 months, α-syn aggregation was increased in all regions and was seen in numerous cell-body-like structures (**Fig. 2a**). PFF-CD163KO females had significantly greater MJF14+ staining in the piriform cortex and amygdala, and a similar trend in SN (vs. PFF-WT female **Fig. 2b-d**). Likewise, pSer129 α-syn immunostaining revealed pathology in all PFF-injected animals (vs. MONO) in both contralateral and ipsilateral striatum after 6 months (**Supplem. Fig. 5e-f**). This was substantially higher in CD163KO mice and particularly enriched in CD163KO females across structures: striatum, piriform cortex and amygdala (**Table 1, Fig.2e-g** and **Supplem. Fig. 5e-j**) (vs. PFF-CD163KO males, PFF-WT females, and MONO-CD163KO females). Notably, PFF-CD163KO females showed more α-syn pathology in the contralateral hemisphere than PFF-WT females in all regions, and this was also true for PFF-CD163KO males in the amygdala (**Supplem. Fig. 5e, g, i**). No positive staining for any of the antibodies was found in PBS-injected mice and none or rare structures were found in α-syn MONO-injected animals (**Supplem. Fig. 5k-n**). This suggests a role for CD163 in the brain’s ability to avoid/handle α-syn pathology, with a particular relevance in females.

**Fig. 2.**
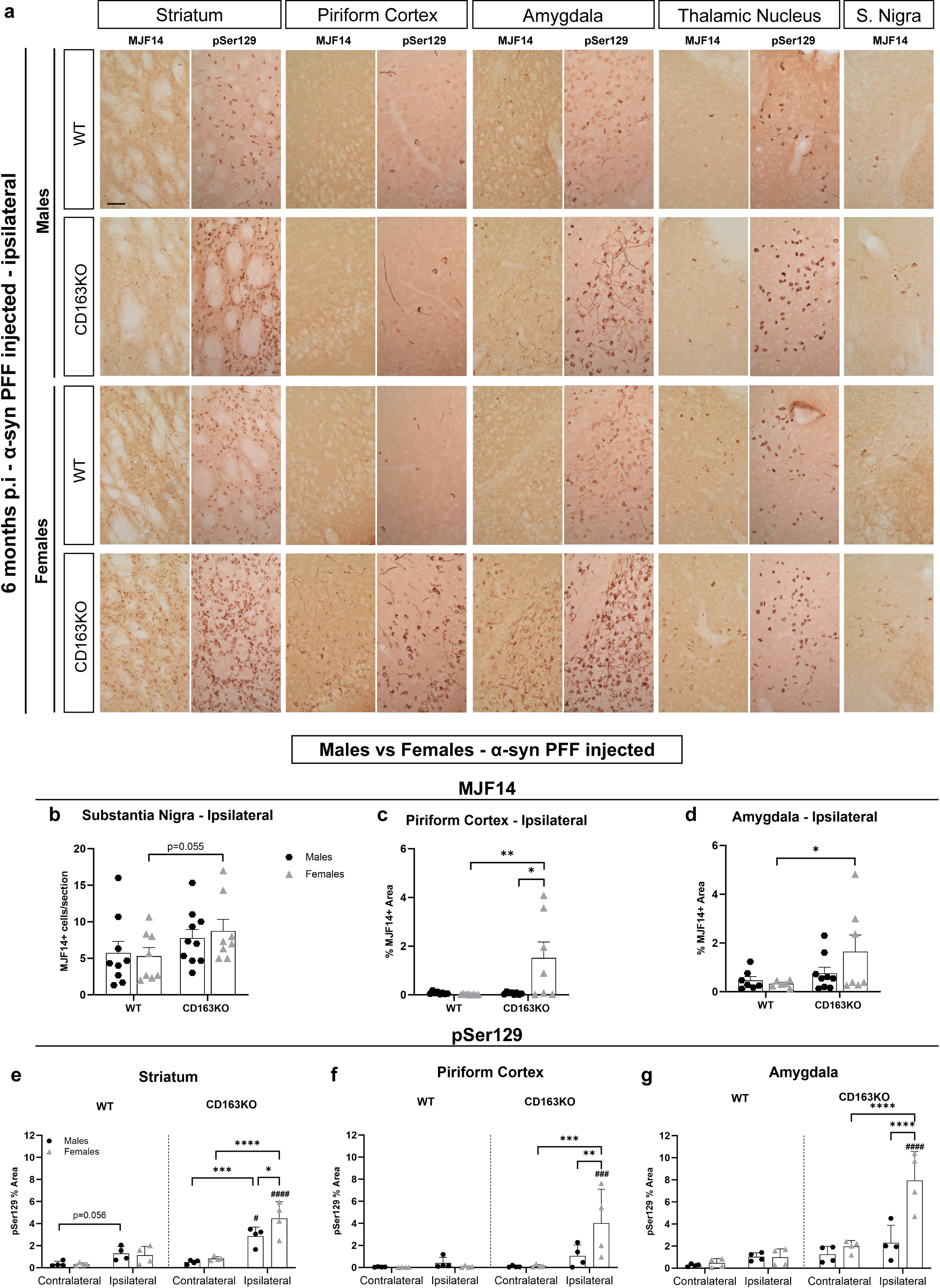
Aggregated and phosphorylated α-syn expression in the striatum and interconnected regions after α-syn PFF injection. **a)** Representative images of MJF14 (left columns) and pSer129 α-syn immunostaining (right columns) in α-syn PFF animals 6-months p.i. **b)** Bar graphs with individual points show the total number of cells per section in the SN, and the percentage of area covered by MJF14+ staining in the **c)** ipsilateral piriform cortex and **d)** ipsilateral amygdala, or pSer129+ staining in the **e)** striatum, **f)** piriform cortex and **g)** amygdala, 6-months post-α-syn-PFF injection. Scale: 50μm applies to all. Values are Mean±SEM (n=4-10). Statistics: Two-way ANOVA followed by Sidak’s multiple comparison test. P values are given according to the number of symbols e.g. (*p<0.05, **<0.01, ***<0.001, ****<0.0001). # different to the corresponding WT group.

### α-syn aggregation is associated with p62 accumulation

As α-syn aggregation has been associated with the accumulation of the autophagy-related protein p62/SQSTM1^33^, we performed immunohistochemistry against p62 (**Fig. 3**). We observed increased p62+ structures in the ipsilateral SN 1-month p.i. in all PFF-injected animals (vs. MONO). This was milder in PFF-CD163KO females, whose increase failed to reach significance (p=0.07) and held significantly fewer p62+ structures when compared to CD163KO males and WT females (**Table 1**, **Fig. 3a-b**). However, at 6-months, while p62+ aggregates in the SN decreased in most of the PFF-injected animals, this was not seen in the PFF-CD163KO females that tend to have higher p62 signal than WT females (p=0.053) (**Fig. 3d-e**), suggesting a delayed change in p62 accumulation in this group. Similarly PFF-CD163KO females showed significantly more p62 staining in the piriform cortex and amygdala structures (vs. CD163KO males and WT-females) (**Table 1**, **Fig. 3g-h, j-k**). p62 expression was associated with α-syn pathology, since the number of p62-expressing cells moderately correlated with the MJF14+ cell numbers in the SN (at 1 and 6 months) and strongly with pSer129 pathology in the piriform cortex and amygdala at 6 months in PFF-injected animals (**Fig. 3c, f, i, l**). Indeed, confocal analysis of p62 and MJF14 in the different brain areas at 6 months revealed that some, but not all, p62 structures co-localized with aggregated α-syn (particularly cytoplasmic perinuclear aggregates of half-moon shape) (**Supplem. Fig. 6**). No p62 co-localization was seen with Iba1+ microglia (**Supplem. Fig. 7i**) or GFAP+ astrocytes (**Supplem. Fig. 8h**); and no p62+ aggregates were found in PBS and MONO-injected animals (**Fig. 3m**). However, confocal imaging confirmed the presence of p62+ structures in TH+ dopaminergic neurons (**Supplem. Fig. 9a, arrowheads**). Altogether, our data suggest that the lack of CD163 results in a temporal delay of the α-syn-induced changes of p62-expression in females, but not in males.

**Fig. 3.**
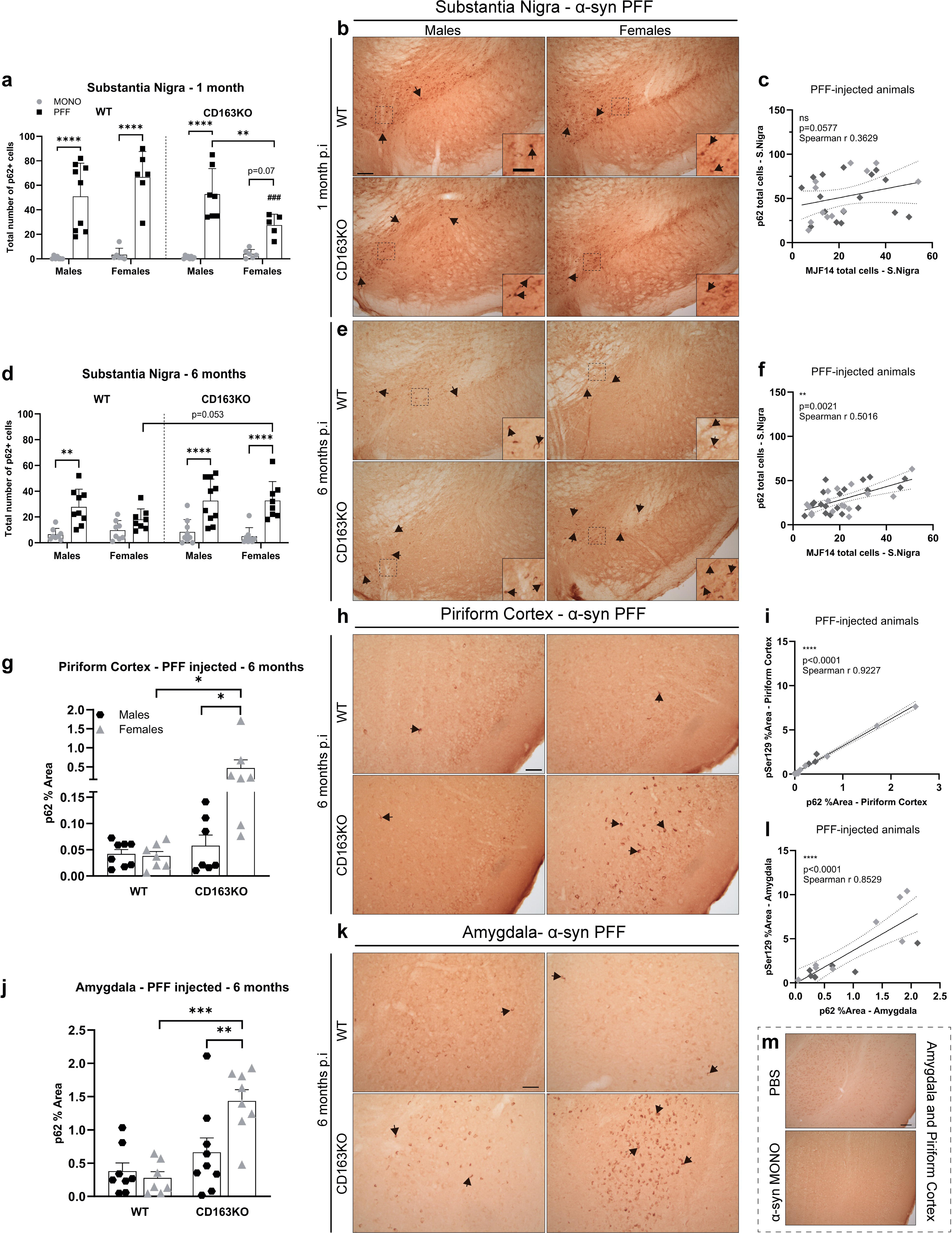
Differential expression of p62/SQSTM1 autophagy marker in the Substantia Nigra, Piriform Cortex and Amygdala after α-syn PFF injection. Bar graphs with individual points illustrate the total number of p62+ cells in the SN at **a)** 1- and **d)** 6-months p.i. **b)** Representative images of p62/SQSTM1 immunostaining in α-syn PFF animals at **b)** 1- and **e)** 6 months p.i. in the SN, **h)** piriform cortex and **k)** amygdala. **g)** Bar graphs with individual points illustrate the percentage of area covered by p62+ staining 6 months after α-syn PFF-injection in WT/CD163KO males and females in the piriform cortex and **j)** amygdala. The number of p62+ cells correlated to MJF14+ cells in the SN at **c)** 1- and **f)** 6-months p.i. The percentage of area covered by p62+ staining correlated to the one covered by pSer129+ staining **i)** in the piriform cortex and **l)** amygdala at 6-months p.i. in all α-syn PFF animals. In the correlation plots: light grey symbols represent females and dark grey males. Scale bar represents 100μm and 50 μm in cropped images in **b,e)** and 50 μm in **h,k)**. Values are Mean±SEM (n=6-10). Statistics: Spearman two-tail p-values (*<0.05, **<0.01, ***<0.001, ****<0.0001), Spearman r and best-fit slope with 95% confidence intervals are plotted. Two-way ANOVA followed by Sidak’s multiple comparison test. *p<0.05, **<0.01, ***<0.001, ****<0.0001

### Microglial response to α-syn is influenced by the lack of CD163

To evaluate changes in the immune response due to CD163 deletion, we performed immunohistochemistry for different immune-relevant proteins. CD68 expression was used as an indirect measurement of immune phagocytic activity and was analyzed in the striatum, piriform cortex, amygdala, and SN (**Fig. 4a-r, Supplem. Fig. 7a-h**). At 1 month, CD68 expression in PBS-injected animals was very low and similar across hemispheres and groups in any of the areas investigated, except in the ipsilateral SN, which was increased (**Fig. 4d, i, n, Supplem. Fig. 7a**). This suggests that the surgery itself is not sufficient to induce overt phagocytic activity in myeloid cells in the brain, while it also indicates a higher sensibility of the immune cells in the SN to respond to mild-innocuous changes in anatomically connected areas. In general, CD68 expression in MONO-injected brains was bilaterally higher than in PBS-injected animals, although this was not always significant (**Fig. 4 d-r, Supplem. Fig. 7a-e**). Nevertheless, α-syn MONO injections led to a significant ipsilateral CD68 upregulation in the striatum irrespective of sex and genotype (**Fig. 4e,h**), which was also seen in WT males in the piriform cortex (**Fig. 4j**). Therefore, an excess of soluble α-syn is sufficient to induce phagocytic activity in myeloid cells in the brain.

**Fig. 4.**
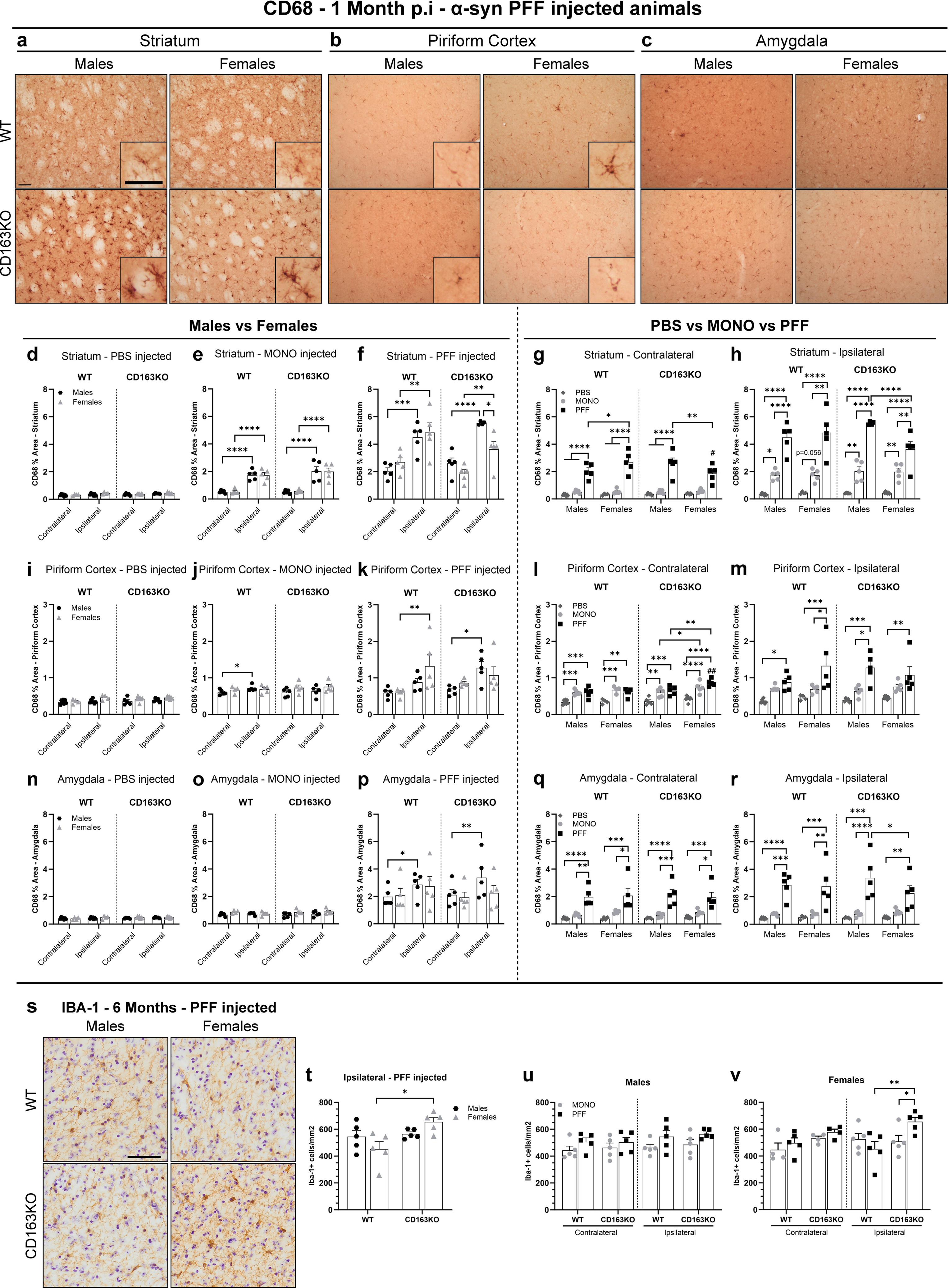
CD163KO females display dysfunctional microglial responses to α-syn PFF injection. **a)** Representative images of CD68 immunostaining in α-syn PFF animals in the striatum, **b)** piriform cortex and **c)** amygdala at 1 month p.i. Scale: 50μm applies to all. Bar graphs with individual points illustrate the percentage of area covered by CD68+ staining after PBS, α-syn MONO and PFF injection in the **d-h)** striatum, **i-m)** piriform cortex and **n-r)** amygdala at 1 month p.i. **s)** Representative ipsilateral images of Iba-1 immunostaining in α-syn PFF-injected animals 6 months p.i. Scale: 50μm applies to all in s). **t)** Bar graphs with individual points show the number of Iba-1 positive cells per mm^2^ in the ipsilateral SN, 6 months after α-syn PFF injection. **u-v)** and in the contralateral and ipsilateral SN, 6 months after α-syn MONO and PFF injection. Values are Mean±SEM (n=5). Statistics: Two-way ANOVA followed by Sidak’s multiple comparison test, or **m)** with Bonferroni corrections applied (i.e. *p<0.0125). P values are given according to the number of symbols e.g. (*p<0.05, **<0.01, ***<0.001, ****<0.0001). # different to the corresponding WT group.

In contrast, PFF-injected animals robustly expressed CD68 in all areas examined (vs. PBS) (**Fig. 4a-r**, **Supplem. Fig. 7c-e**). PFF-induced CD68 upregulation was higher than the one seen in MONO-injected animals (**Fig. 4g-h, m, q-r**), and also enhanced in the contralateral side in all areas (vs. PBS) (**Fig.4 g, l, q, Supplem. Fig. 7d)**. However, CD68 upregulation was greater in the ipsilateral side (vs. contralateral) in all PFF-injected animals in the striatum (**Fig. 4f**), in the WT females and CD163KO males in the piriform cortex (**Fig. 4k**) and only in males in amygdala and SN (**Fig. 4p, Supplem. Fig. 7c)**. This highlights the relevance of the sex in the immune response to α-syn PFF, with an apparent higher sensibility in males. Notably, the PFF-induced CD68 upregulation was significantly reduced in the ipsilateral striatum and amygdala of CD163KO females when compared to CD163KO males (**Table 1**, **Fig. 4 h, r**). Interestingly, in the striatum, both CD163KO males and females showed CD68+ cells of bigger size and shorter ramifications, suggestive of a hypertrophic/phagocytic phenotype in mice lacking CD163 (**Fig. 4a**). These observations suggest that females lacking CD163 portray a reduced CD68-phagocytic response to pathological α-syn. The CD68 phagocytic activity was associated with α-syn aggregation, as supported by the significant positive correlations between MJF14 pathology and CD68 expression in the SN (Spearman r= 0.561, p=0.01), piriform cortex (r=0.7549, p=0.0007) and amygdala (r=0.679, p=0.001).

At 6 months, striatal CD68 expression in PFF-injected animals decreased to those observed in the MONO groups (**Supplem. Fig. 7f**). At this time-point, quantification of Iba1+ microglia in SN (**Fig. 4s-v**) showed an ipsilateral microglia proliferation in PFF-CD163KO females (vs. ipsilateral in PFF-WT and MONO-CD163KO females, **Fig. 4s-t, v**). No significant differences in the number of Iba1+ microglia were found in the SN of the male groups, although ipsilateral numbers were slightly higher (**Fig. 4u**). However, we observed a decrease in the percentage of Iba-1+ type A cells (homeostatic-surveilling) in the ipsilateral side of all PFF-injected animals except in the PFF-CD163KO females (**Supplem. Fig. 7j**). Altogether our results suggest a dysfunctional, long-lasting microglia response to α-syn PFF injection in CD163KO females.

### Differential astroglial responses in α-syn PFF-injected CD163KO females

Due to the relevant role of astrocytes in the immune response in the CNS ^34^, we evaluated the GFAP+ astrocytic compartment in the striatum, piriform cortex, amygdala and SN (**Fig. 5, Supplem. Fig.8 a-e**). At 1-month, intrastriatal PBS injection increased the area covered by GFAP staining in the ipsilateral striatum of all mice (vs. contralateral) (**Fig. 5d**), in CD163KO females in the piriform cortex (**Fig. 5i**) and in WT males and in all females in the amygdala (**Fig. 5n**). This suggests an early response of astrocytes to the surgery that appeared more relevant in females. α-syn MONO injections also increased GFAP expression in the ipsilateral striatum and SN in all mice (**Fig. 5e, Supplem. Fig. 8b**), in piriform cortex of WT (**Fig. 5j**) and only in females in the amygdala (**Fig. 5o**). Interestingly, MONO-CD163KO females showed higher GFAP expression than MONO-CD163KO males in the piriform cortex and amygdala (**Fig. 5 l, m, o, q**). This again suggests a glial response to the monomeric α-syn, as well as to the sex-dimorphism in such response.

**Fig. 5.**
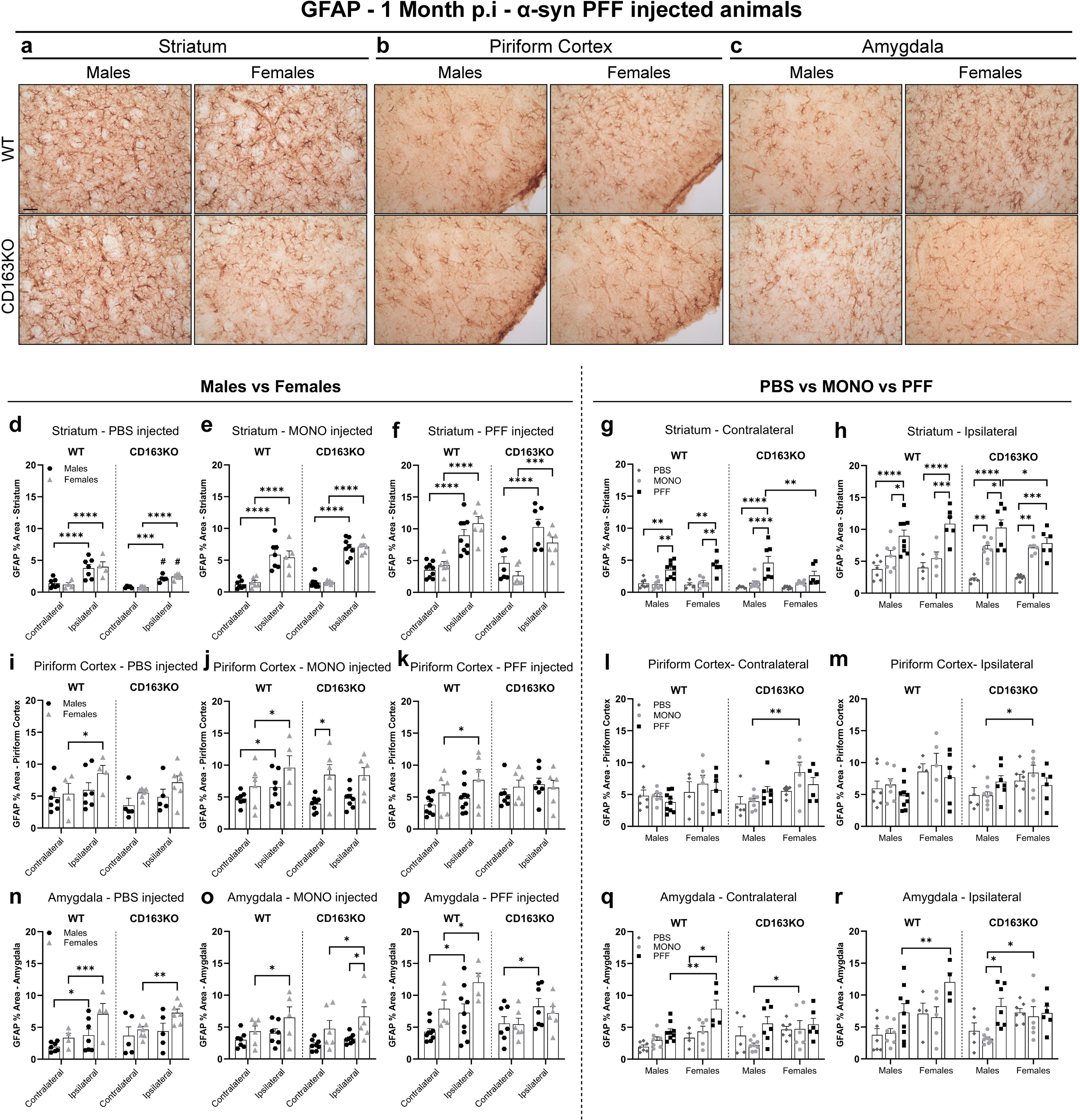
CD163KO females display dysfunctional astrocytic responses to α-syn PFF injection. **a)** Representative images of GFAP immunostaining in α-syn PFF animals in the striatum, **b)** piriform cortex and **c)** amygdala at 1 month p.i. Scale: 50μm applies to all. Bar graphs with individual points illustrate the percentage of area covered by GFAP+ staining after PBS, α-syn MONO and PFF injection in the **d-h)** striatum, **i-m)** piriform cortex and **n-r)** amygdala at 1 month p.i. Values are Mean±SEM (n=5-10). Statistics: Two-way ANOVA followed by Sidak’s multiple comparison test. P values are given according to the number of symbols e.g. (*p<0.05, **<0.01, ***<0.001, ****<0.0001). # different to the corresponding WT group.

The α-syn PFF injections resulted in ipsilateral (vs. contralateral) GFAP upregulation in all animals in the striatum and SN (**Fig. 5f and Supplem. Fig. 8c**), and in all but CD163KO females in the amygdala (**Fig. 5p**), while only WT females showed ipsilateral GFAP upregulation in the piriform cortex (**Fig. 5k**). As observed for CD68, PFF-CD163KO females revealed diminished levels of GFAP expression compared to CD163KO males in the ipsilateral striatum (**Fig. 5h**). The contralateral striatum of all PFF-injected animals, except the CD163KO females, also showed GFAP upregulation (vs. PBS and MONO), but still to a lower extent than in the ipsilateral side (**Fig. 5g).** α-syn PFF injections lead to bilateral GFAP upregulation in the amygdala in WT females, that showed higher GFAP expression than PFF-WT males in both sides (**Table 1**, **Fig. 5q-r**). These observations highlight not only the sex relevance, but also the anatomical differences in the astrocytic response to α-syn. At 6 months p.i., GFAP expression levels in striatum and SN were similar between MONO and PFF animals and across hemispheres (**Supplem. Fig. 8f-g**). In conclusion, our results suggest that CD163 deletion leads to an impaired astrocytic response to α-syn PFF injection in females early in the disease process.

### Robust MHCII upregulation in α-syn PFF-injected males

Expression of major histocompatibility complex II (MHCII) is a widely used marker of responsive microglia, and has been associated with α-syn pathology in rodent models^5^ and in *postmortem* PD brains^3^. As this model with time shows bilateral α-syn pathology, both hemispheres were analyzed. Here, we found that MHCII expression, rarely found in microglia of the healthy brain (as confirmed in the PBS-injected animals (**Fig.6h**)), was upregulated early in those regions with MJF14/pSer129+ α-syn pathology (**Fig. 6a**). In most brain regions, MHCII+ cells exhibited a ramified morphology (likely microglia), however, in the piriform cortex, we only observed numerous elongated rod-shape MHCII+ cells, resembling perivascular macrophages (**Fig. 6a)**. At 1-month, we observed substantially more MHCII+ ramified cells in the ipsilateral SN of all α-syn PFF-injected males (vs. contralateral), but mainly in CD163KO males that revealed more MHCII expression than MONO-CD163KO males (**Fig. 6b)**. Interestingly, fewer cells expressed MHCII in the female groups, and only the PFF-CD163KO females tend to show an ipsilateral MHCII upregulation (p=0.06) (**Fig. 6b**). The robust ipsilateral MHCII upregulation in the CD163KO males, was also seen in the striatum, where MHCII expression was significantly higher than its contralateral (**Fig. 6c**), but also greater than MONO-CD163KO males and PFF-CD163KO females, which showed low levels of MHCII expression (**Table 1**, **Fig. 6d**). After 6-months MHCII expressing-cells were rare in the SN of all animals (**Fig. 6e**). In the ipsilateral striatum, despite the very low number of MHCII+ cells, PFF-CD163KO females showed significantly more MHCII+ ramified cells than PFF-WT females (**Fig. 6f**). No differences were found in the ipsilateral amygdala (**Fig. 6g**). Therefore, males, predominantly CD163KO, respond strongly to α-syn by upregulating MHCII early, while females do not. However, the absence of CD163 in females resulted in a late MHCII+ response to α-syn pathology. This suggests a temporal delay and a sex-dependent role for CD163 in the antigen presentation response to α-syn by innate immune cells.

**Fig. 6.**
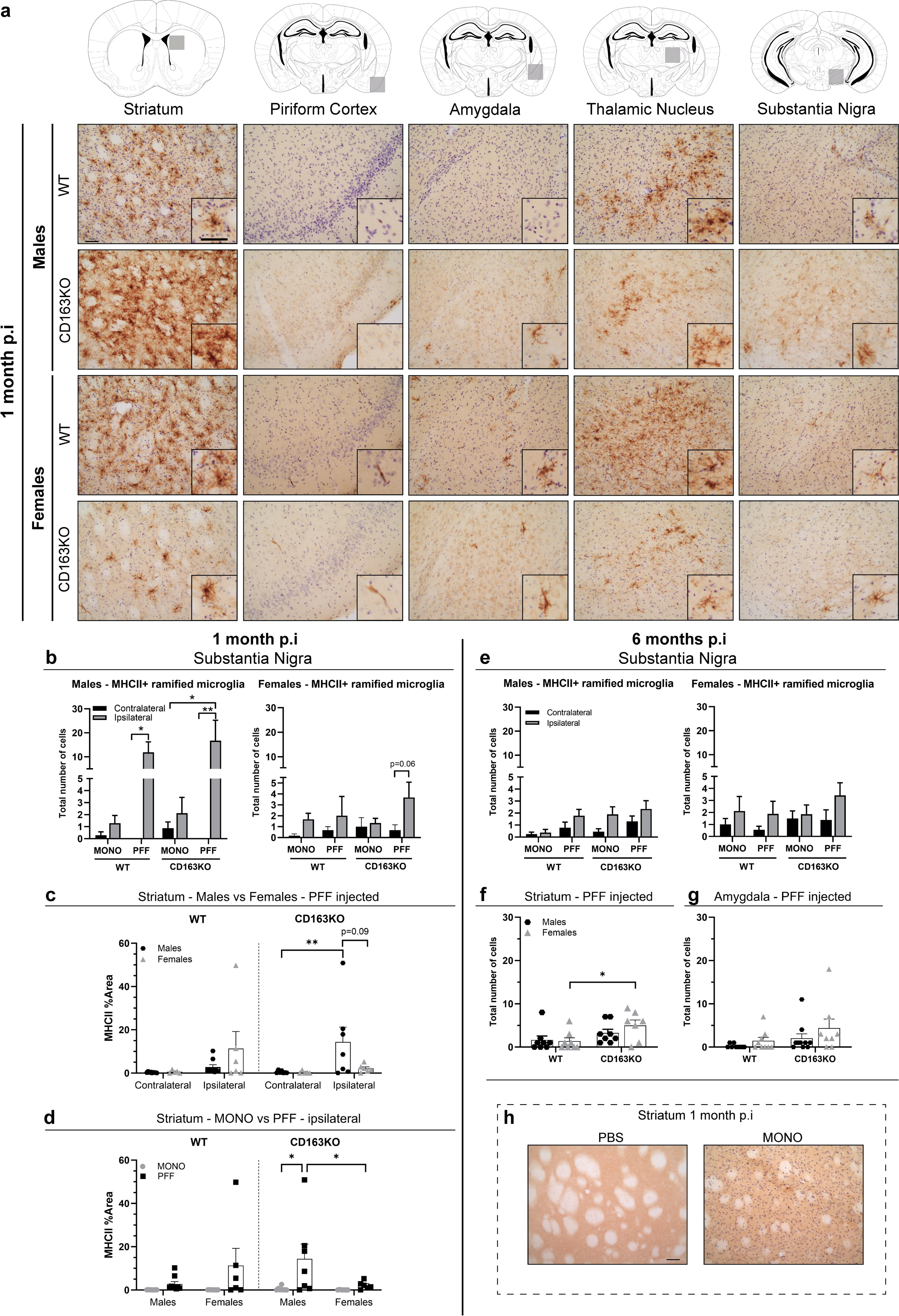
Expression of MHCII in the Striatum and interconnected brain regions after α-syn PFF injection. **a)** Representative images of MHCII immunostaining in α-syn PFF animals 1 month p.i. Areas marked by gray squares in the atlas sections above represent areas photographed. Bar graphs illustrate the number of MHCII+ cells in the SN at **b)** 1- and **e)** 6 months p.i. **c-d)** Bar graphs with individual points show the percentage of area covered by MHCII in the striatum of at 1 month p.i. **f)** Bar graph with individual points show the number of MHCII+ cells in the striatum and **g)** amygdala at 6 months p.i. **h)** Representative images of MHCII immunostaining in PBS and α-syn MONO-injected animals. Scale bar: 50μm applies to all. Values are Mean±SEM (n=6-10). Statistics:c) Kruskal-Wallis test; b, d-e) Two-way ANOVA followed by Sidak’s multiple comparison test. *p<0.05, **p<0.01.

### α-syn PFF CD163KO mice show a sex dimorphic T cell response

Changes within the innate immune response will ultimately influence the adaptive immune system. α-syn aggregates and pathology have been shown to lead to MHC system upregulation that in turn will activate T cells ^35,36^. Additionally, since both CD4 and CD8 T cells have been observed in *postmortem* PD brains, we wondered if CD163 deletion would enhance T-cell brain infiltration in the striatum (**Fig. 7**). Interestingly, at 1-month p.i, we observed a significant bilateral CD4 T helper cell infiltration in WT animals following α-syn MONO injection, although significantly higher in the ipsilateral side (**Fig. 7b, d, e**). Yet, this was not seen in CD163KO animals (**Fig. 7b, d**). We also found low, although significant, ipsilateral CD8 T cell infiltration with α-syn MONO injection in WT females and CD163KO males (**Fig. 7f**).

**Fig. 7.**
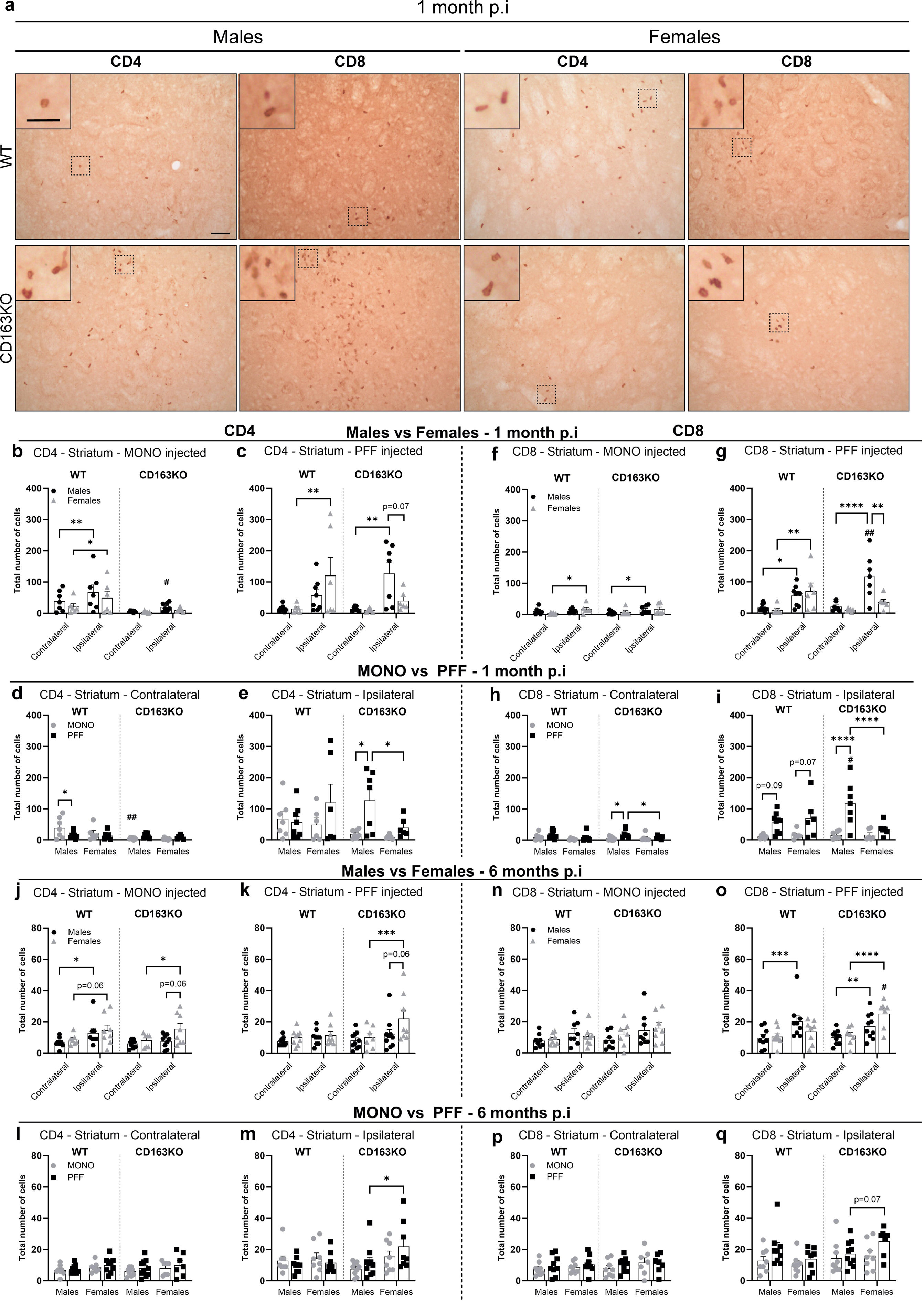
CD4-T and CD8-T cell infiltration in the Striatum after α-syn PFF injection. **a)** Representative images of CD4 and CD8 immunostaining in α-syn PFF animals at 1 month p.i. Scale bars represent 50μm and 25μm for cropped images. Bar graphs with individual points show the total number of **b-e, j-m)** CD4+ and **f-I, n-q)** CD8+ cells after α-syn MONO and PFF injection at **b-i)** 1 month and **j-q)** 6 months p.i. Values are Mean±SEM (n=6-10). Statistics: Two-way ANOVA followed by Sidak’s multiple comparison test. P values are given according to the number of symbols e.g. (*p<0.05, **<0.01, ***<0.001, ****<0.0001). # different to the corresponding WT group.

α-syn PFF-injections led to increased CD4 T cells infiltration exclusively in the ipsilateral striatum in WT females and CD163KO males (vs. contralateral). However, this increase was not seen in PFF-CD163KO females, which had significantly fewer CD4 T cells than CD163KO males (**Table 1**, **Fig.7c, e**). Similarly, α-syn PFF injection led to significant CD8 T-cell infiltration in the ipsilateral striatum of all animals except CD163KO females. CD8 T cell infiltration was, however, enhanced in CD163KO males, which also show contralateral CD8 T cell infiltration (vs. MONO) even though to a lower degree than in the ipsilateral striatum (**Fig.7f-i**). Therefore, MONO α-syn injections lead to prompt bilateral CD4 T cell infiltration in WT but not in CD63KO animals, while PFF α-syn injections resulted in ipsilateral CD4 and CD8 T cell infiltration in all mice, though of higher magnitude in CD163KO males and lower in CD163KO females.

After 6 months, despite the overall decrease in T cell numbers, α-syn MONO injected mice showed an ipsilateral increase in CD4 T cells in all animals except CD163KO males (vs. contralateral) (**Fig. 7j**). However, in the α-syn PFF-mice, only CD163KO females had significantly higher CD4 in the ipsilateral striatum (vs. contralateral) and than in PFF-CD163KO males (**Fig.7 k, m**). At this time point, CD8 infiltration was observed in the ipsilateral striatum of PFF-injected males, both WT and CD163KO, but also in CD163KO females, which showed more CD8+ cells than WT females and a similar trend vs. CD163KO males (p=0.07) (**Fig. 7 o, q**). Hence, the lack of CD163 resulted in a significant long-term T-cell infiltration, which was otherwise absent in WT females. As a result, CD163 seems to play a role in modulating the innate immune responses and in turn adaptive immune responses to α-syn PFF injection, in a sex-specific manner.

### α-syn PFF-CD163KO females showed greater long-term dopaminergic loss

Given our observations on the alteration of α-syn pathology and sex-specific dysfunctional immune responses in CD163KO, we assessed whether such events led to distinct dopaminergic neurodegeneration (**Fig. 8**). Densitometry analysis of striatal TH+ fibers revealed a 18-21% decrease in the ipsilateral side in all α-syn PFF groups at 1 month (vs. PBS and MONO) (**Fig. 8a-b**), which remained after 6-months, in α-syn PFF males (**Fig. 8c-d**). However, PFF-CD163KO females showed a further reduction to 44%, resulting on significantly lower axonal TH-density than MONO-CD163KO, PFF-WT females and PFF-CD163KO males (**Table 1**, **Fig. 8e-f, h**). No rostro/caudal pattern of dopaminergic denervation was seen (**Fig. 8d, f**).

**Fig. 8.**
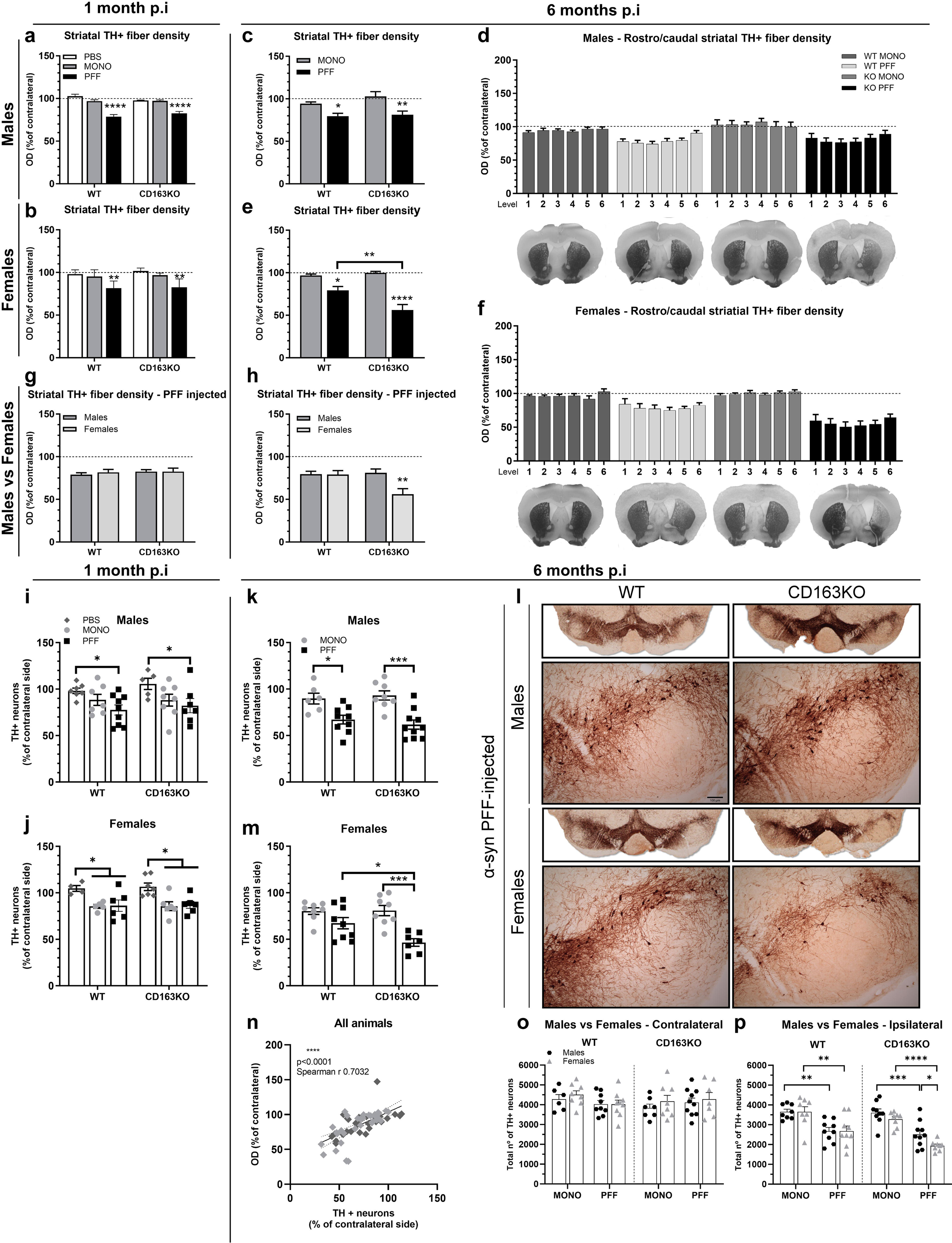
Striatal axonal density and stereological quantification of TH+ cell bodies in the Substantia Nigra. **a-h)** Bar graphs illustrate the semi-quantitative measurement of TH+ axonal fiber density expressed as the percentage of the contralateral striatal side, post-PBS, α-syn MONO and PFF injection in the striatum. Average values of the 6 rostro-caudal striatal levels quantified are shown in **a)** Males and **b)** Females at 1 month p.i. Average and separate values of the 6 rostro-caudal striatal levels quantified are shown in **c-d)** for Males and **e-f)** Females at 6 months p.i, together with representative photos of one section at striatal level 0.38 from bregma in each group. Average values of the 6 rostro-caudal striatal levels quantified in α-syn PFF-injected animals are shown at **g)** 1 month p.i and **b)** 6 months p.i. **l)** Low magnification photos show representative nigral sections immunostained for TH and higher magnification photos show the ipsilateral SN from WT and CD163KO Male and Female mice after α-syn PFF injection. **i-p)** Stereological quantification of TH+ neurons in SN (**i-j, k, m**) as percentage of contralateral or **o-p)** total number of neurons. Bar graphs with individual points illustrate the average number of surviving TH+ neurons in PBS, α-syn MONO and PFF-injected WT and CD163KO males and females at **i-j)** 1- and **k,m)** and α-syn MONO and PFF-injected animals at 6-months p.i; **n)** Correlation between the striatal TH+ axonal density and the number of TH+ SN neurons (light grey symbols represent female and dark grey males). **o)** Quantification of total number of TH+ neurons in the contralateral SN and in **n)** ipsilateral SN. Scale bar for lower magnification photos represent 12,5mm, and in higher magnification photos 100μm as shown in **l)**. Values are Mean±SEM (n=4-10). Statistics: Two-way ANOVA followed by Sidak’s multiple comparison test. *p<0.05, **<0.01, ***<0.001, ****<0.0001.

Stereological quantification of SN TH+ cell bodies showed decrease of TH+ neurons (as % of contralateral) in PFF-injected males and females (vs. PBS) but also MONO-injected females (**Fig. 8i-j**). However, after 6 months, PFF-WT and PFF-CD163KO males showed a 33% and 38% loss of dopaminergic neurons, respectively, in the ipsilateral SN (vs. MONO) (**Fig. 8k**). Notably, PFF-CD163KO females showed the biggest ipsilateral loss of dopaminergic neurons (54%) compared to MONO-CD163KO and PFF-WT females (**Fig. 8m, l**). No difference in the total TH+ numbers in the contralateral SN was seen (**Fig. 8o**), however, we found significant neuronal loss in the ipsilateral SN of all PFF-injected animals after 6 months. This was particularly enhanced in PFF-CD163KO females, which showed significantly lower number of surviving neurons compared to MONO-CD163KO females, and PFF-CD163KO males (**Table 1**, **Fig. 8p**). As expected, the striatal axonal TH+ density correlated significantly to the TH+ neuronal number in the SN (**Fig. 8n**). Moreover, the number of p62+ cells negatively correlated with the percentage of surviving TH+ neurons in SN of α-syn PFF animals (**Supplem. Fig. 9b-h**). This correlation was stronger in males (**Supplem. Fig. 9c**) and particularly driven by CD163 deletion (**Supplem. Fig. 9e**), suggesting that p62 accumulation is a significant factor in driving neurodegeneration in CD163-deficient males. These findings suggest a sex-dependent neuroprotective role for the CD163 receptor in the progressive neurodegenerative process triggered by α-syn aggregation in SN.

### CD163 deletion alters the gene expression profile of macrophages and microglia after α-syn PFF injection

Based on the literature at 2 months p.i, the immune response related to the α-syn injection in the model has been resolved, α-syn pathology is significant, but no TH neuronal loss has occurred ^37,38^. Therefore, to comprehend the consequences of CD163 absence in the early immune response associated with neuronal α-syn pathology prior to cell death, we performed SMART-seq2 at 2 months p.i. in the females. This was performed in two FACS-sorted brain immune populations expressing: CD45 low/intermediate/CD11b^+^, the Microglia, and CD45 high/CD11b^+^ (macrophages (brain-border associated and infiltrated), neutrophils and NK cells), which we refer to as the Macrophage population (**Fig. 9, Supplem. Fig. 10**).

**Fig.9.**
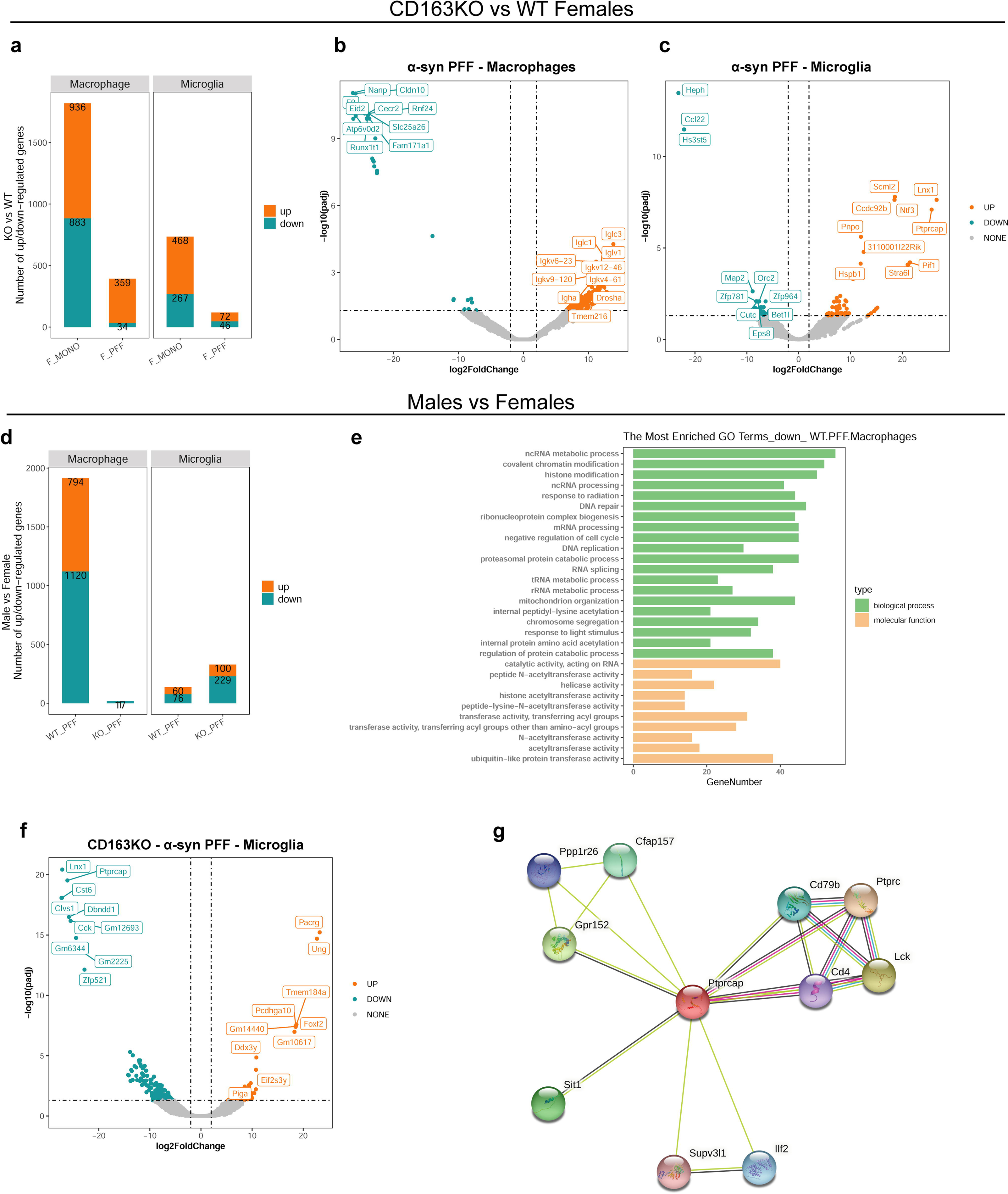
Microglial *Ptprcap* upregulation in CD163KO Females and transcriptomic sex-differences. **a)** Bar graph showing the number of up/downregulated genes in CD163KO vs. WT females in Macrophage and Microglia populations 2 months p.i. **b)** Volcano scatter-plots (-log10(padj) *vs* log2FoldChange) of DEGs (CD163KO vs. WT) in Females-PFF Macrophages and **c)** microglia. **d)** Bar graph showing the number of up/downregulated genes in Males vs. Females (α-syn PFF groups) in Macrophage and Microglia populations 2 months p.i. **e)** Gene Ontology (GO) analysis on the downregulated genes in WT-PFF (Males vs. Females) macrophages. **f)** Volcano scatter-plot (-log10(padj) *vs* log2FoldChange) of DEGs (Males vs. Females) in CD163KO-PFF Microglia. **g)** STRING protein-protein interaction network with focus on *Ptprcap* gene. Statistics: abs (logFC)>1, FDR<0.05.

Differentially expressed gene (DEGs) analysis revealed a reduced ability of the macrophage population to respond to α-syn PFF injection (vs. MONO) in CD163KO females compared to WT. This was paralleled by a differential response of the microglia population to α-syn pathology (**Supplem. Fig. 10a**). In the two populations, very few or no DEGs were commonly shared between WT and CD163KO females, suggesting that CD163 deficiency drastically affected the immune response to the α-syn pathology (**Supplem. Fig. 10b**).Within the macrophage population, the most upregulated genes in PFF-WT females were related to DNA damage response and regulation of catabolic processes (*Cecr2, Fzd6*), while genes associated with amino acid/protein metabolism (*Mpst, Akr1c12, Rsc1a1, Zfp354c*) and cytoskeleton organization (*Frmd5, Fhod3*) were downregulated (**Supplem. Fig. 10c**). CD163KO females showed a drastically reduced macrophage response to α-syn PFF, suggesting an impaired ability to mount an otherwise “normal” response to the aggregated protein. Enriched gene ontology (GO) functional annotations showed that the downregulated genes in the CD163KO females were involved in regulation of calcium homeostasis and metabolism (**Supplem. Fig. 10d, g**), known to play a central role in immune cell activation ^39^. The microglia population of the PFF-WT females showed upregulation of genes related to: peptide delivery to MHCI (*Lrmp*) and regulation of TLR4 surface expression (*Cnpy4*); while genes associated with: vesicle coating (*Clvs1*), DNA repair (*Mcm8, Trib2*) and metabolic mechanisms (*Got1l1, Acot11, Kyat1*) were downregulated (**Supplem. Fig. 10e**). Interestingly, microglia from CD163KO females revealed the most significant DEGs (lowest p value and highest fold change (PFF vs. MONO)). Here, we saw upregulation of *Lnx1*, a gene related with ubiquitination and activation of proteosomal degradation mechanisms, *St8sia1,* associated with sphingolipid metabolism, and *Ptprcap,* encoding a key regulator of CD45-CD4 T-cell interaction through the TNFR1 pathway. Genes related to DNA damage response (*Atad5, Smad9*), apoptosis modulation (*Dab2ip*), cell migration (*Dnai3*) and cytoskeleton organization (*Slain1*) were downregulated (**Supplem. Fig. 10f**).

Since CD163KO females showed a significant increase in α-syn-mediated neurotoxicity, we further focused on this group and assessed the response to PFF and MONO α-syn in CD163KO vs. WT females. The number of DEGs in PFF-CD163KO females (vs. PFF-WT) macrophages were significantly fewer than those seen when comparing the MONO females (**Fig. 9a**) and were associated with downregulation of genes related to amino acid/protein metabolism (*Nanp*), cytoskeletal dynamics (*Fam171a1*), tight junctions and BBB integrity (*Cldn10*) (**Fig. 9b**). Microglia gene expression was also affected (**Fig. 9a**), with downregulation of iron metabolism-related genes (*Heph*), and heparan sulfate biosynthesis mechanisms (*Hs3st5*). Moreover *Ccl22,* a chemokine related to T-cell migration was also downregulated in PFF-CD163KO females, suggesting that CD163 expression in myeloid cells influences peripheral cell infiltration in females. GO analysis revealed few specific pathways affected by gene downregulation and were limited to ubiquitin binding and DNA catalytic activity pathways (not shown). The most upregulated genes were: *Ntf3,* which encodes Neurotrophin3 a protein involved in neuronal survival and differentiation, and, once again, *Lnx1* and *Ptprcap* (**Fig. 9c**). Altogether, these results suggest that the early transcriptomic immune alterations seen in CD163KO females may contribute to their higher susceptibility to α-syn PFF.

To investigate potential sex-differences, we compared DEGs in Males vs. Females after α-syn PFF injection (**Fig. 9d-f**). Macrophage transcriptomes were very different between sexes in the PPF-WT group, where, in contrast to females, males presented many downregulated genes (1120)(vs. female) mostly related to modulation of epigenetic machinery and DNA repair mechanisms (**Fig. 9e**), confirming a sex-dimorphism in the macrophage response to α-syn PFF. Remarkably, this sex difference in DEGs was lost upon CD163 deletion, which mostly resulted in a few downregulated genes in male macrophages (**Fig. 9d**). However, the CD163 deletion resulted in a bigger number of genes differentially regulated in male microglia (vs. females), also compared to those in the WT group (**Fig. 9d**). The most highly upregulated gene found in PFF-CD163KO male microglia was *Pacrg*, which encodes the Parkin co-regulated protein, a protein found in Lewy bodies ^40^ and related to aggresome formation and increased autophagy ^41^. Confirming our previous observations, the most significant downregulated genes in CD163KO male microglia were *Lnx1* and *Ptprcap*, which, as mentioned, were the most upregulated genes in CD163KO females (vs. WT and MONO) (**Fig.9f**). Interestingly, STRING analysis of the Ptprcap protein shows interaction with CD4, TNFR1 and TWEAK pathways (**Fig. 9g**). These results confirm the sex-dimorphic behavior of CD163 in the α-syn-associated immune response.

### CD163 deletion does not alter the *in vitro* response of macrophages to α-syn PFF

We have previously shown that the CD163KO does not drastically affect the *in vitro* immune response of macrophages ^15^. To evaluate if this is also true for the α-syn PFF-induced response, we isolated BMDM from WT and CD163KO mice and stimulated them with either IL-10 or IFNγ for 24h prior to α-syn PFF treatment (**Supplem. Fig.11**). Expression of CXCL10, TNF and iNOS after BMDM priming with IFNγ, were significantly enhanced by α-syn PFF treatment in both WT and CD163KO males (**Supplem.Fig.11a, c-d**) and females (**Supplem.Fig.11e, g-h**). IL-1β expression, was significantly increased in all WT and CD163KO males BMDM primed with IFNγ and incubated with α-syn PFF (**Supplem.Fig.11b, f**); but not in CD163KO females BMDM (**Supplem.Fig.11f, red bars**), suggesting an irregular response to α-syn PFF. VCAM-1 and VEGFR1 (*Ftl-1*) mRNA were also measured, but no differences between groups were seen (not shown). Thus, the absence of CD163 does not drastically affect the ability of BMDM to acutely respond to extracellular α-syn PFF.

## DISCUSSION

To investigate the involvement of the CD163 receptor in the α-syn induced neurodegeneration and associated immune response, we used the α-syn PFF PD model in CD163KO mice. Based on prior observations regarding CD163 in PD patients, and the sex dimorphism observed in immune system and PD, sexes were analyzed separately^42,43^. To study the progressive nature of the response 1-, 2- and 6-months survival were used as previously ^37,38,44^. α-syn PFF injection resulted in early and sustained motor alterations in CD163KO males, but not in females. However, α-syn PFF injection led to a progressive α-syn pathological aggregation and p62 accumulation significantly higher in CD163KO females after 6 months. α-syn PFF-injections resulted in early immune response, with upregulation of CD68, MHCII, T-cell infiltration and astrogliosis, that was diminished or absent in CD163KO females. Nevertheless, this group revealed a delayed but significant immune response at 6 months as shown by the Iba1+ microglia proliferation in SN, and the highest MHCII expression and T-cell infiltration in the striatum. Remarkably, this differential immune response and the increased pathological α-syn in CD163KO females culminated in higher nigrostriatal dopaminergic degeneration. RNA-seq analysis at 2-months p.i. revealed that CD163KO female macrophages lack the ability to respond to α-syn PFF. Additionally, CD163 deletion in females induced an altered microglial phenotype with upregulation of genes related to pro-inflammatory pathways, which, together with changes in the intracellular milieu, may contribute to the enhanced α-syn PFF-induced neurodegeneration observed in CD163KO females. Overall, our data support a novel and sex-dimorphic role for CD163 in the α-syn-induced immune response and neurodegeneration.

Intra-striatal injection of murine α-syn PFF induced aggregation and Ser129 phosphorylation of the endogenous α-syn in the striatum and in anatomically connected areas as before ^35,45,46^. This was prominent in CD163KO animals and particularly enhanced in CD163KO females. Thus, CD163 deletion potentiates α-syn phosphorylation, increasing toxic α-syn aggregates^32,47^, particularly in females. α-syn aggregation and neuronal toxicity is associated with autophagic dysfunction and p62 accumulation ^33^. Accordingly, PFF-injected mice showed p62 accumulation that paralleled the α-syn pathology; and thus, although delayed compared to the other groups, it was higher in CD163KO females at 6 months. Thus, CD163 absence resulted in a more prominent long-term autophagic changes in females. p62 colocalized with MJF14+ α-syn aggregates, and it was observed in SN dopaminergic neurons in agreement with our prior observations in the rat PPF PD model ^30^. Interestingly, we have recent *in vitro* observations showing that sCD163, produced during monocyte pro-inflammatory activation, increases the uptake of extracellular α-syn by monocytes and microglia, which should in turn clear it ^26^. Our results here suggest that CD163 deficiency may compromise the ability of myeloid cells to properly respond to α-syn, consequently promoting its pathological intraneuronal aggregation and p62 accumulation, particularly in females.

PD-related inflammation involves both brain and peripheral immune cells ^2^. Accordingly, the α-syn-PFF PD model shows microgliosis ^30,48^, presence of CD163+ cells in the brain^20^ and changes in peripheral immune cells ^49–51^. We observed CD68 upregulation suggestive of an early phagocytic microglia activation, which coincided with MHCII upregulation. This MHCII response was higher in the CD163KO males, suggesting an enhanced early adaptive immune activation, as confirmed by the higher CD4 and CD8 T-cell infiltration. In contrast, this early immune response was not seen in the PFF-CD163KO females, which also lacked the GFAP upregulation seen in all other PFF-groups. However, after 6 months, when this immune response was resolved in most groups, it became relevant in the PFF-CD163KO females, which showed a significant unique SN microgliosis (Iba-1+ proliferation), higher MHCII expression, and persistent T cell infiltration in the striatum. This might ultimately contribute to higher neurodegeneration as T-cells seem to mediate dopaminergic loss in PD models ^52,53^. In conclusion, CD163KO females showed an inability to timely respond to early α-syn-associated pathological events, which resulted in a delayed, but long-lasting neuroinflammation, involving both innate and adaptive cells.

Site and sex-specific changes were seen throughout the study. Although pathology was seen in most of the anatomically connected regions to the striatum, the amygdala was one of the areas showing the most changes, as before ^54^. Notably, the amygdala in *postmortem* PD showed significant immune changes and α-syn pathology ^55^. Here, the astrocytes in the amygdala seemed to be the most responsive (reacting to mild conditions such as PBS or MONO), and was highly affected by the CD163 deletion, showing more bilateral pSer129 pathology at long-term. However, microglia in the SN seemed particularly sensitive, with CD68 upregulation even after striatal PBS injections. This suggests a differential susceptibility to α-syn-related pathology in different brain areas, as seen in PD brains, and an anatomically distinctive glia response to brain changes. This is in agreement with the anatomical differences in microglia in healthy brains ^56^ and in an α-syn PD model ^57^. We also saw sex-specific changes (Table 1); in general, male mice (WT and CD163KO) tend to show stronger changes than females after PFF α-syn injections: with higher early upregulation of MHCII in SN, higher CD68 expression in the amygdala and long term CD8 T cell ipsilateral infiltration in the striatum. On the other hand, GFAP upregulation seemed more consistent in females, while MHCII expression was lower. Additionally, WT females showed less robust p62 expression in SN 6 months after PFF α-syn injections. This agrees with a previous report of a more “neuroprotective” profile of microglia in females^58^ vs. males, with a more inflammatory response and higher MHCII expressors^58,59^. Of relevance to the disease modeling, MONO α-syn injections were not innocuous and showed significant CD68 and GFAP upregulation, together with CD4 T-cell infiltration at short and long term in WT animals. Interestingly this MONO response was affected by CD163 absence as no early T cell infiltration was seen in the CD163KO mice, and only females CD163KO showed it at the long term, which further supports a role for the CD163 in the adaptive response. The MONO α-syn injections resulted in a small bilateral decrease of TH+ cells in SN after 1month, which might be associated with the CD4 T-cell infiltration observed.

Further corroborating a differential immune response, our transcriptomic data showed that CD163KO leads to an impaired macrophage response and a differential microglia response to α-syn PFF in females. Most DEG in the female CD163KO macrophages were downregulated and involved processes such as calcium homeostasis regulation, which plays a central role in immune cell activation ^39^. Also, others related to cytoskeleton organization, BBB integrity and immune cell transmigration, which could be associated with the early decreased T-cell infiltration in this group. α-syn PFF WT female microglia showed upregulated genes coding for proteins previously related to α-syn immune response (MHC system and TLR4) ^2^. However, microglia from CD163KO females showed downregulation of *Ccl22,* a chemokine involved in the regulation of leukocyte migration ^60^ that might also contribute to the decreased early T-cell infiltration. This may suggest that CD163 expression is involved in the early peripheral cell infiltration.

Of particular interest in CD163KO females, we observed upregulation of *Ptprcap,* a key regulator of CD45-CD4 T-cell interaction and activation ^61^, also associated with activation of the TNFR1 and TWEAK pathways. *Ptprcap* upregulation has been previously shown in the midbrain of the α-syn viral-vector PD model^62^. Interestingly, sCD163 is suggested to act as a decoy receptor for TWEAK, regulating TWEAK-induced pro-inflammatory canonical NF-κB activation ^63^. Notably, TWEAK has been associated with dopaminergic cell death and the pro-inflammatory activation of astrocytes in the MPTP PD model ^64,65^; and TWEAK is increased in PD patients’ serum ^66^. Therefore, upregulation of *Ptprcap* and possible TWEAK-mediated inflammation may be also involved in the higher susceptibility of CD163KO females to α-syn PFF. Although, further studies are required to investigate this possible connection.

The higher burden of α-syn pathology and the differential immune response observed in the α-syn PFF CD163KO females resulted in increased dopaminergic loss. CD163+ cells are found in the brain in rodent PD models ^19,20,30^ and in PD *postmortem* brains^67^. We previously reported increased CD163+ cell numbers and CD163 expression levels (in both sexes) on blood monocytes from patients with early PD^11^. Our data here supports a protective role for the CD163 receptor on myeloid cells. This is agreement with our recent observations in REM sleep behavior disorder patients (prodromal PD), where higher numbers of CD163 cells in the blood were associated with lower immune activation in the SN and better dopaminergic transmission in the putamen as shown by positron emission tomography ^27^. Intriguingly, we observed here that CD163 neuroprotective role seems especially relevant in females. Remarkably, we have previously seen that shedding of CD163 was only increased in the sera from female PD patients, but not in males, supporting a sex-difference regarding the CD163-system/cells during PD ^26^. Further suggesting a sex dimorphism in the PD-associated immune response, blood monocytes transcriptome differs between male and female PD patients ^10^. Our histological data and RNA-seq analysis of the innate cells in the α-syn PFF-WT animals further corroborate a sex-dimorphic immune response, which may have consequences for the neuronal health. Sex differences in immune cells have been described before ^58,68,69^. The observed differential behavior of the CD163 and dimorphic immune response might be related to the higher PD risk in males ^70^ and the differential presentation of the disease among sexes ^9^. Our data suggest that CD163 is also involved in this sex-dimorphism, as its deletion in females resulted in a defective ability of immune cells to properly respond to the initial α-syn pathological effects, ultimately influencing the long-term degenerative outcome; while in males the absence of CD163 increased motor defects. Our histology data suggests anatomical differences in the immune response that might be diluted and missed by using the whole brain for transcriptomics. Novel spatial transcriptomic techniques might resolve these questions in the future.

Our behavioral analysis supports a relevant role for the immune system in the symptomatic disease presentation. While all α-syn injected mice show behavioral changes at 1 month (vs. PBS) this was more pronounced in the PFF CD163KO male, but not females. Moreover, only PFF CD163KO male showed asymmetry at 6 months p.i.. This also points towards a sex-difference in the symptomatic manifestation of motor defects, ultimately influenced by the immune environment. The CD163KO has been associated with distinct temporal influences on neurobehavioral tests and mortality in a mouse model of intracerebral hemorrhage ^16^. However, this study included only males and shorter time-points. CD163KO strongly enhanced the disease severity and led to imbalanced CD4 Th1/Th2 responses in a model of rheumatoid arthritis ^15^. Interestingly, despite the α-syn pathology and the dopaminergic degeneration seen in WT, we only observed long-term behavioral changes in the CD163KO mice, thus supporting a role for the immune environment in motor performance. We and others have previously reported that the α-syn PFF rodent PD model in WT (male or female) does not exhibit robust motor defects despite the nigral degeneration ^30,45,71,72^. However, changes in the challenging beam in pesticide exposed PFF-PD model have been seen only in males, and this was not related to bigger nigral TH loss or α-syn pathology, but to dopamine turnover and inflammatory changes^72^. A predominant affection in males has been also seen in other PD relevant studies in mice ^73,74^, and male MPTP-intoxicated monkeys showed higher inflammation markers than females ^75^. Such observation suggests that the immune response significantly influences the motor phenotype particularly in males. This might be exerted by soluble cytokines ^76,77^ or peripheral cell infiltration, which can potentially influence microglia and neuronal function/firing ^78^. Accordingly, sickness behavior was associated with peripheral inflammation responsible for sensorimotor defects ^79^. Together with the observed early increase in MHCII expression and the T cell infiltration, the behavior impairments in CD163KO males could also be partially contributed by the neuronal α-syn pathology and autophagic changes besides the dopaminergic neurodegeneration.

Overall, our results suggest that, although CD163 deletion does not affect the *in vitro* acute monocytic response to fibrillar α-syn-induced activation, *in vivo* it results in a differential early transcriptomic innate immune response. In males, CD163 deletion leads to early increased T-cell infiltration and motor impairment, while in females it may contribute to their higher susceptibility to α-syn PFF and promote increased long-term neurodegeneration. This might be related to changes in the border associated macrophages, which are CD163+ and recently proposed to be crucial mediators of the T-cell infiltration and neurodegeneration in an α-syn PD model ^80^. Interestingly, CD163 expression is described in the disease associated microglia (DAM) in AD ^25^. According to scRNA-seq studies, CD163 is absent in homeostatic microglia ^21–23^, thus its expression in the DAM indicates an ectopic expression, or as suggested before, a loss of identity of the microglia during disease by downregulating its characteristic markers while gaining others ^81,82^, or alternatively to a de-differentiation of infiltrated macrophages. Notably, a recent scRNA-seq study in the brain from PD patients also highlighted CD163 expression in the DAM ^24^. In AD, the subpopulation of CD163+ amyloid-responsive microglia were depleted in AD cases with APOE and TREM2 risk variants, also suggesting a protective role ^83^. Furthermore, CD163 upregulation has been also associated with macrophages with neuroprotective capacity in an AD model ^84^. Thus, it could be speculated that CD163 upregulation is associated with a protective compensatory mechanism exerted by myeloid cells occurring both in the brain and in the blood of PD patients ^11,24^. In conclusion, we demonstrate for the first time that the α-syn-associated innate immune responses involving CD163 expression differ between males and females, suggesting that the sex-dimorphism in PD may be due to immune differences. Therefore, future designs of immunomodulatory treatments should account for distinct immune phenotypes. In addition, our data suggest that the CD163 upregulation in the myeloid compartment during neurodegeneration is a compensatory neuroprotective mechanism. Future studies may in further molecular detail explain the role of CD163 in brain diseases, as well the mechanisms defining the gender dimorphism observed.

## ACKNOWLEDGEMENTS

We are grateful to the FACS core facility for the technical help with the FACSAria III high speed cell sorter and the Imaging core facility for technical support with the Upright Widefield Slide Scanner microscope. Funding support for the research covered in this article was provided by the Bjarne Saxhof Fund administered through the Danish Parkinson’s Foundation (MRR), the Novo Nordisk Foundation (DEO, NNF17OC0028806, MRR) and Independent Research Fund Denmark (9039-00217B, MRR). The Lundbeck Foundation grants R223-2015-4222 and R248-2016-2518 for Danish Research Institute of Translational Neuroscience-DANDRITE (PHJ & CB). SAF was a recipient of a PhD fellowship from the Graduate School at the Health Faculty, Aarhus University when performing the experiments described here.

## AUTHORS CONTRIBUTION

SAF, ZV and MRR did stereotaxic surgeries; SAF, IK, ZV and AK performed and analyzed behavior studies and histological samples. SAF and GUT performed histological experiments, SAF performed microglia/macrophage isolation, selection and RNA purification; RKR performed qPCR analysis; CL and YL analyzed the SMARTseq2 data. AE, PS, and SKM have contributed to the CD163KO mouse line studies. CB and PHJ produced the recombinant mouse α-syn PFF. MRR conceived the idea, SAF and MRR designed the study and wrote jointly the manuscript. All authors contributed to the manuscript and approved the final version.

## SUPPLEMENTARY MATERIAL

Supplementary material is available online.

## DATA AVAILABILITY

The authors confirm that the data supporting the findings of this study are available within the article and its supplementary material, or otherwise are available from the corresponding author, upon reasonable request.

