## Supplementary info for "Sex-dimorphic neuroprotective effect of CD163 in an α-synuclein mouse model of Parkinson’s disease"

#### Supplementary information

##### Materials and Methods

##### Animals

Adult male and female CD163KO (CD163$^{tm1.1(KOMP)Vlcg}$) mice and WT (C57BL/6N) littermates (3-4 months old) (n=179 total (n=95 males (50 WT + 45 CD163KO); n=84 females (43 WT+ 41 CD163KO)) were used in this study [1]. Mice weighted 20-25g (12 weeks old) at the time of the surgery and were housed a maximum 4-6 per cage, with *ad libitum* access to food and water, in a climate-controlled facility under a 12h/12h night/daylight cycle. All animal experiments were approved and performed under humane conditions in accordance with the ethical guidelines established by the Danish Animal Inspectorate and following EU legislation.

##### Experimental design

To generate a PD model, murine alpha-synuclein pre-formed fibrils (α-syn PFF) were used due to their higher seeding efficiency in mice compared to human α-syn PFF [2,3]. In parallel, phosphate buffered saline (PBS) was used as an absolute control, and murine monomeric α-syn (MONO) was used as a control for α-syn pathology, based on its reported lack of seeding capacity *in vitro* [4] and *in vivo* [5]. Mice received unilateral intrastriatal injection of PBS, α-syn MONO, or α-syn PFF and were euthanized 1 or 6 months post-surgery. (**Supplementary Fig. 1**). The 1-month group included: PBS (Males WT n=7 & CD163KO n=5; Females WT n=4 & CD163KO n=7), α-syn MONO (Males WT n=7 & CD163KO n =8; Females WT n=6 & CD163KO n=6), and α-syn PFF (Males WT n=9 &CD163KO n =7; Females WT n=6 &CD163KO n=6). While the 6-months group consists of: α-syn MONO (Males WT n=8 & CD163KO n=9; Females WT n=8 &CD163KO n=8), and in the α-syn PFF (Males WT n=9 & CD163KO n=10, Females WT n=9, & CD163KO n=8). Motor behavior was assessed on the week before each end-point using the Challenging Beam Test and the Cylinder test. Mice were euthanized, perfused and brains processed for immunohistochemical staining. At times, not all animals of each group and time were used for each immunohistochemical marker to secure enough tissue to analyze multiple proteins. Figures show individual points, to inform on the number of animals analyzed for each marker or alternatively n number is given in figure legends.

For whole population RNA sequencing, a third and fourth group of mice (batch 1 and batch 2) received bilateral intrastriatal injection of α-syn MONO (Males WT n=7 & CD163KO n=4*; Females WT n=7 &CD163KO n=5), and α-syn PFF (Males WT n=7, & CD163KO n =5; Females WT n=7 & CD163KO n=5). Triplicates were used for sequencing. Mice were sacrificed 2 months post-surgery, their brains dissected and the immune cells (microglia & macrophages) isolated and FACS sorted for RNA purification and subsequent SMART-seq2 sequencing. *CD163KO-MONO males were excluded from the analysis due to inconsistencies in the technical replicates.

For the *in vitro* experiment, bone-marrow derived macrophages (BMDM) were isolated from CD163KO and WT C57BL/6N male and female mice, aged 6-12 weeks (n=5 per group).

**Protein purification and aggregation of mouse α-synuclein**

Murine α-syn was recombinantly expressed in E. coli BL21 DE3 bacteria using the pRK172 plasmid encoding for mouse α-syn (kind gift from Prof. Virginia Lee (University of Pennsylvania)). The bacteria were pelleted by centrifugation and resuspended in buffer (50 mM Tris, 1 mM EDTA, 0.1 mM DTE, 0.1mM PMSF, pH 7.0). The suspension was lyzed by sonication in a Branson Sonifier (Output control 7; dutycycle 50), and centrifuged at 20,000 *x g* for 30 minutes at 4°C. The supernatant was boiled for 5 min and centrifuged at 48.000 *x g* for 30 min at 4° C. Preceding ion exchange purification, the supernatant was dialyzed in 20 mM Tris pH 6.5, and filtered through a 45μm filter. Murine α-syn was purified on POROS HQ 50 ion exchange chromatography with a continuous gradient of 0% - 100% 2 M NaCl in 20 mM Tris pH 6.5. The fractions with murine α-syn were isolated and further purified by reverse phase chromatography (C18) in order to remove nucleotides, and lipids bound to α-syn. This step also removes endotoxins (lipoglycans) from the sample (<0.5 EU/mg, confirmed using the Pierce™ Chromogenic

Endotoxin Quant Kit, ThermoScientific™). The pure protein was dialyzed in 20 mM ammonium bicarbonate, lyophilized, and stored at -20°C. In order to produce sterile PFF, mouse α-syn was solubilized in PBS (7.2 mM $Na_2HPO_4$, 2.8 mM $NaH_2PO_4$, 140 mM NaCl, pH 7.4) to a final concentration of 7 mg/mL, and further sterile filtered through a 0.22 µm sterile filter in a LAF bench. The solution was allowed to aggregate at 37°C, 1000 RPM for 10 days. The insoluble α-syn PFF were isolated from unbound α-syn MONO by 20,000 *x g* centrifugation for 30 min at 20° C and further resuspended in fresh sterile PBS. Protein concentration was determined by Bicinchoninic acid protein concentration assay (Pierce).

**Recombinant α-syn fibrils sonication and stereotaxic surgery**

On the day of surgery, α-syn PFF were sonicated for 40 minutes (70ms off/30ms on, 30% output power, Branson 250 sonifier), and their hydrodynamic average length was determined prior and post-surgery by Dynamic Light Scattering (DLS) (Wyatt DynaPro NanoStar), at 25°C. Fibrils were of average =33.87nm (range: 25.90-41.10nm), thus within the optimal size range of 29-49nm previously reported [6], and remained so during the surgery day (not shown). Mice were anesthetized (Medetomidine hydrochloride (1mg/ml), Midazolam (5mg/ml), and Fentanyl (0,05mg/ml) in 0.9% NaCl for a final i/p. dose of 20ml/kg) and placed in a stereotaxic apparatus (Stoeling). α-syn PFF or α-syn MONO (2µl of 5 µg/µl, in sterile PBS) were unilaterally (or bilaterally for RNA analysis) injected into the striatum: AP +0.7/+0.9mm (females and males respectively), ML +/- 2.0mm (from bregma), and DV -3.0mm (from dura) [7]. A s.c. antagonist solution (Flumazenil (0.1 mg/ml), Naloxone (0.4mg/ml), and Antipamezol hydrochloride (5mg/ml) in 0.9% NaCl for a final dose of 20ml/kg) was used and fully awaken animals were placed back in their cages and i.p injected with Buprenorphine (0.3mg/mL) for pain-relief.

**Behavioral Tests**

The Challenging Beam Test was performed using a beam with four 25 cm frames of progressively decreasing width as described before [8]. Animals were trained for 2 days to transverse the beam from the widest (frame 1) to the narrowest section (frame 4). On the test day, a mesh grid was placed over the beam leaving approximately 1cm space between the grid and the beam surface. Animals were video-recorded while crossing the grid-surface with five trials per animal. Frame 1 was excluded from the analysis due to variable reactions after first contact with the beam.

Videotapes of each animal were rated for time to traverse the beam, number of errors, and number of steps for each of the trials and averaged per animal.

The spontaneous activity on the Cylinder test was assessed as formerly [8]. Spontaneous activity in a glass cylinder was videotaped for 3-7 minutes and a test was considered valid if the number of rears were equal or > 20. To evaluate motor dexterity, the number of hindlimb steps were counted. For vertical activity, the number of rears were counted. Hindlimb steps were counted when the animal placed a paw in a different position on the cylinder floor, toward a frontal or backwards movement. The percentage of total contralateral and ipsilateral steps, contralateral paw use, and total hindlimb steps were calculated.

**Perfusion and Tissue processing**

Animals were euthanized with an overdose of pentobarbital (400mg/mL, 1:10 I.P.). Upon respiratory arrest, animals were transcardially perfused through the ascending aorta with ice-cold 0.9% NaCl solution, followed by 4% paraformaldehyde (PFA in 0.1M phosphate buffer, pH 7.4). PFA-perfused brains were removed and post-fixed in the same 4% PFA solution for 2h and transferred to a 25% sucrose solution (in 0.02M NaPBS) overnight for cryoprotection. Brains were subsequently sliced into 40-µm-thick coronal sections using an HM 450 Slicer Microtome (Brock and Michelsen, Thermo Fisher Scientific) and separated in series of 6 for the Striatum and of 4 for the SN. Sections were stored at -20°C in an anti-freeze solution.

**Immunohistochemistry with DAB detection and Immunofluorescence**

Immunohistochemical staining was done on free-floating brain sections as previously described [9]. Briefly, sections were blocked in appropriate 5% normal serum and then labeled overnight with anti-tyrosine hydroxylase (TH, 1:750, Millipore), anti-aggregated α-syn MJFR-14-6-4-2 (MJF14,1:25000, Abcam), anti-phosphorylated α-syn (pSer129 α-syn 1:3000, Cell Signaling Technology), MHCII (1:400, eBioscience), p62/SQSTM1 (1:2000, Nordic Biosite), CD68 (1:1000, BioRad), Iba-1 (1:1000, Wako Fujifilm), GFAP (1:5000, Abcam), CD4 (1:500, BD Biosciences) or CD8 (1:500, eBioscience) in 2.5% serum and 0.25% Triton-X-100 in KPBS at room temperature. Sections were washed with KPBS, pre-blocked for 10min in 1% serum and 0.25% Triton-X-100, and incubated with the appropriate biotinylated secondary antibodies (1:200, Burlingame, CA Vector Laboratories) for 2 hours. Afterward, avidin-biotin-peroxidase complex

(ABS Elite, Vector Laboratories) was used and visualized using 3,3-diaminobenzidine (DAB) as a chromogen with 0.01-0.1% $H_2O_2$. Sections were mounted on chrome-alum gelatin-coated slides and coverslipped. For MHCII and Iba-1 staining, sections were counterstained with Cresyl violet (0.5% solution). Slides were analyzed using a Leica DMI600B brightfield microscope unless specified.

For immunofluorescence, free-floating sections were blocked in the appropriate 5% normal serum and then labeled overnight with anti-TH (1:750, Millipore), anti-aggregated α-syn MJFR-14-6-4-2 (1:10000, Abcam), p62/SQSTM1 (C-terminus) (1:1000, Nordic Biosite), GFAP (1:5000, Abcam) and Iba-1 (1:1000, Wako Fujifilm). Sections were washed with KPBS and incubated for 2 hours with species-specific fluorochrome-conjugated secondary antibodies (Alexa-Fluor 488 or 647, Invitrogen) plus DAPI (1:2000, Sigma-Aldrich A/S) for nuclear staining. Sections were mounted on chrome-alum gelatin-coated slides with Dako fluorescent mounting medium. Confocal images were obtained using a LSM 710 Meta Confocal microscope (Zeiss) with a 20X/0.8 M27 objective. Extraction of single z-frame and maximum intensity projections were performed with ImageJ (Fiji) software.

**Densitometric analysis of dopaminergic axonal striatal innervation**

Striatal density of TH+ dopaminergic fibers were measured by analysis of optical density at 6 different rostro-caudal levels [7]: AP: +1.10; +0.62; +0.38; +0.14; -0.22; -0.58 mm relative to bregma. Immunostained sections were scanned using a densitometer (EPSON Perfection 3200 (1800 dpi resolution, grayscale), and the digital images acquired were analyzed with ImageJ (Fiji) software using a grayscale. The optical density in each section was corrected to an unspecific background measured in the corpus callosum of the same section. Data are shown as a percentage of the ipsilateral side vs. the contralateral side.

**Stereology and microscopic analysis**

Unbiased stereological estimation of total TH+ cells in the SN was performed by an observer blind to the animal's identity, using the optical fractionator principle [10,11] and a Bright field Leica DM600B microscope with the NEWcast program (Visiopharm). A 1.25X low power objective (HCX PL Fluotar, Germany) was used to outline the SN based on its anatomical landmarks. Dopaminergic TH+ neurons were counted with a 40X objective (Leica, Germany) in a series of 1:4 sections covering the full SN (8-10 nigral sections per animal) from the rostral corner of the pars compacta to the caudal end of the pars reticulata (located between -2.70mm and -3.88mm from bregma [7]) with a counting frame of 56.89µm x 42.66 µm and a step length of 110-165 as to count a minimum of 100 cells per SN and a CE<0.1.

Using a Bright field Leica DM600B microscope, the total number of cells with aggregated MJF14+ α-syn, and p62+ structures were manually counted in SN sections (three equally distant per staining; between -2.46 and -3.64 mm from bregma) [7]. Total number of MHCII+ cells in the SN were manually counted in a series of 6 equally distant midbrain coronal sections (-2.46 to -3.88 mm from bregma), and CD4+ and CD8+ T cells were counted in striatal sections (three equally distant per staining; 1.18 and -0.22mm from bregma). At 6 months p.i, MHCII+ cells in the striatum and amygdala were manually quantified in three equally distant coronal sections (1.18 and -0.22mm from bregma).

Enhanced Focal Images (EFI) captured using the Upright Widefield Slide Scanner microscope (UWSSM) were used to analyze the area covered by immunostaining in two-three coronal sections stained for: phosphorylated α-syn (pSer129) in the striatum (between 1.18 and -0.22mm from bregma), amygdala and piriform cortex (0.26 and -2.18 mm); MJF14+ aggregates in the amygdala and piriform cortex (0.26 and -2.18 mm); p62-expressing cells in the amygdala and piriform cortex (-0.26 and -2.18mm); CD68 in the striatum (1.18 and -0.22mm) and in the SN (-2.70 and -3.88mm); GFAP in the striatum (1.18 and -0.22mm) and in the SN (-2.70 and -3.88mm), and MHCII in the striatum (1.18 and -0.22mm from bregma), using ImageJ (Fiji).

Iba-1+ cells were counted with a 40X objective in 3 equally distant SN sections (-2.70mm and -3.88mm) aided by the VIS module (Visiopharm program). The counting frame (56.89µm x 42.66 µm) was randomly located by the VIS module and methodically moved to sample the entire delineated region of the SN (step length of 75 µm) and all cells inside the counting frame were counted. A positive cell had a cresyl violet-stained nucleus covering a Iba1+ cell body. Four cellular profiles were defined as previously described [12]. The number of Iba-1+ cells per $mm^2$ was calculated according to the area sampled in each section (calculated by the VIS module) and the total number of cells counted per section. The percentage of each morphological cell type was calculated as the total % of A,B,C, and D type in each section and averaged per animal.

**Brain cells isolation and fluorescence activated cell sorting for RNA isolation**

Animals were transcardially perfused through the ascending aorta with ice-cold PBS, brains were extracted and cells isolated using the Adult Brain Dissociation Kit (Miltenyi Biotec) in accordance with the manufacturer's protocol. Briefly, brains were cut into approximately 0.5 cm pieces and transferred into gentleMACS C Tubes containing the provided enzymes. Tissue was dissociated using the gentleMACS Octo Dissociator with Heaters for 30 min in the appropriate program. Homogenates were filtered through 70µm cell strainers and the cell suspension was centrifuged at 300xg, 10 minutes at 4°C. Cells were resuspended in a Debris Removal Solution (in DPBS containing $CaCl_2$, $MgCl_2$, 1g/L D-Glucose, and 36mg/L Pyruvate, Gibco) and centrifuged at 4°C and 3000xg for 10 minutes. Gradient centrifugation formed three phases and the top two layers containing myelin debris were removed. Cells were washed with 1x PBS and counted in the MOXI Z Mini Automated cell counter (ORFLO).

Freshly isolated brain cells were blocked with CD16/CD32 (clone 2.4G2, Mouse BD Fc block, BD Pharming) and 10% goat serum in PBS for 10min at 4°C. Single-cell suspensions were incubated in darkness at 4°C for 30 min with CD11b antibody conjugated to BV421 dye (50µg/mL, clone M1/70, BioLegend) and CD45 antibody conjugated to APC dye (0.2mg/mL, clone 30-F11, BioLegend). Cells were washed with PBS and 0.5% BSA, centrifuged at 4°C and 400xg for 5 minutes, and resuspended in cold PBS. Propidium iodide (PI) was added to the cell suspension for identification of dead cells. Microglia and macrophage populations were sorted into separate tubes containing PBS, in the FACSAria III high speed cell sorter (BD Biosciences, San Jose, CA) according to CD11b (405nm laser and 450/40 bandpass filter) and CD45 (633nm laser and 660/20 bandpass filter) (**Supplementary Fig. 2**). Once sorted, microglia (200000-500000 cells) and macrophage (10000-30000 cells) cell suspensions (>90% purity after sorting) were centrifuged at 4°C and 400xg for 5 minutes, resuspended in RLT Plus Lysis buffer (QIAGEN, Germany) with 1% β-Mercaptoethanol and vortexed for 1 minute for cell lysis. The solution was further homogenized using a 1mL syringe and a 21gag needle. Total RNA was extracted using RNeasy® Mini Kit (QIAGEN) according to the manufacturer's protocol.

**SMART-Seq2, gene mapping, expression and analysis**

Total purified RNA was sent to BGI Hong Kong where Switching Mechanism at 5' End of RNA Template (SMART-seq2) sequencing service was requested. Samples were tested for quality control using Agilent 2100 Bio analyzer (Agilent RNA 6000 Nano Kit). RNA integrity (RNI) >7 was considered suitable for sequencing. Gene expression analysis was performed through full population RNA sequencing using the SMART-seq2 method according to BGI's standard methodology.

Reads mapped to rRNA were removed and raw data was obtained. The sequencing reads that contained low-quality, adaptor-polluted, and high content of unknown base (N) reads processed were removed before downstream analyses. Clean reads were mapped to reference using Bowtie2 [13], and then gene expression level was calculated for each sample with RSEM [14], a software package for estimating gene and isoform expression levels from RNA-Seq data. Based on the gene expression level, we identified the DEGs (Differentially expressed genes) between α-syn PFF vs. MONO, Male vs Female, and WT vs. CD163KO groups. We use DESeq2 package of R [15] to detect the DEGs, determining at a cutoff of FDR corrected p-value≤0.05 and |log2(fold change)|≥1 as DEGs.

Principal component analysis (PCA) was performed on the Macrophage and Microglia population respectively based on the rlog of dds result from DESeq2 and draw the diagrams with the ggplot2 function of R. Batch effect was corrected and three outlier samples were removed according to the results of PCA diagrams (**Supplementary Fig. 12**). With DEGs, we performed Gene Ontology (GO) and KEGG pathway classification and functional enrichment using the ClusterProfiler package of R [16]. GO has three ontologies: molecular biological function, cellular component, and biological process. We performed biological process and molecular biological function enrichment respectively, showed when relevant.

##### *In vitro* treatment of bone marrow-derived macrophages and gene expression analysis

Bone marrow derived macrophages (BMDMs) were prepared by flushing the femur and tibia of CD163KO or WT mice, aged 6-12 weeks, with RPMI-1640. Flushed bone marrow was filtered through a 70 µm cell strainer (Corning®), and 6 x $10^6$ cells were plated in uncoated petri dishes (Greiner Bio-one) in RPMI-1640 with 10% Fetal Bovine Serum (FBS, Biowest), penicillin and streptomycin and 2mM L-Glutamine (Gibco), supplemented with 10% L929 conditioned medium. Cells were supplemented with RPMI with 10% FBS, P/S, L-glut, and 10% L929 medium on day 4, and on day 7 the cells were harvested with TrypLE (Gibco) and plated in 24 well plates for downstream assays. Plated BMDMs were incubated with 10 ng/ml IL-10 or IFNγ (Peprotech) for

24 hours and subsequently stimulated with PBS control or 1μM murine α-syn PFF for 6 hours, after which the cells were lyzed in TRIdity G™ (Applichem). RNA was purified using the Direct-zol RNA kit (Zymogen), according to the manufacturer's protocol. The quantity and quality of the RNA was measured using a Nanodrop ONE (Thermo Fischer Scientific), and cDNA was prepared using the High-Capacity cDNA Reverse Transcription Kit (Applied Biosystems). qPCR was performed using the KAPA SYBR® fast low ROX (Sigma Aldrich), 5 ng RNA, and 0,4 μM combined forward/reverse primer on Applied Biosystems 7500 fast qPCR machine, and peptidylprolyl isomerase A was used as reference gene to calculate the relative gene expression ($2^{-\Delta Ct}$).

| Target | GeneID | Sequence (forward/reverse) |
|---|---|---|
| *Ppia* | NM_013494.4 | 5′-ATGGTCAACCCCACCGTG-3' |
| | | 5'-TTCTTGCTGTCTTTGGAACTTTGTC-3' |
| *Cxcl10* | NM_021274.2 | 5'-GGGCCATAGGGAAGCTTGAA-3' |
| | | 5'-GGATTCAGACATCTCTGCTCATCA-3' |
| *Il1b* | NM_008361.3 | 5'-TGGCAACTGTTCCTGAACTCA-3' |
| | | 5'-GGGTCCGTCAACTTCAAAGAAC-3' |
| *Tnf* | NM_013693.3 | 5'-GGGTGATCGGTCCCCAAA-3' |
| | | 5'-TGAGGGTCTGGGCCATAGAA-3' |
| *Nos2* | NM_010927 | 5'-GCCACCAACAATGGCAACAT-3' |
| | | 5'-TCGATGCACAACTGGGTGAA-3' |
| *Flt1* | NM_010228 | 5'-GAGGAGGATGAGGGTGTCTATAGGT-3' |
| | | 5'-GTGATCAGCTCCAGGTTTGACTT-3' |
| *Vcam1* | NM_011693 | 5'-CTCTTACCTGTGCGCTGTGA-3' |
| | | 5'-CTTCAGGGAATGAGTAGACCTCC-3' |

**Supplementary Table 1. Primers used for gene expression analysis of bone marrow-derived macrophages**

#### Statistical analysis

All analysis were done by researchers blind to the group's identity. All statistical analysis were performed using GraphPad Prism v.10.0.1 (GraphPad Software, California, USA). Gaussian Normality was first assessed using normality distribution with a Shapiro-Wilk normality test followed by Two-way ANOVA and, when appropriate, by a post-hoc Sidak's multiple comparisons test. Bonferroni corrections were applied when appropriate. Alpha was set at 5%.

### Supplementary Figures

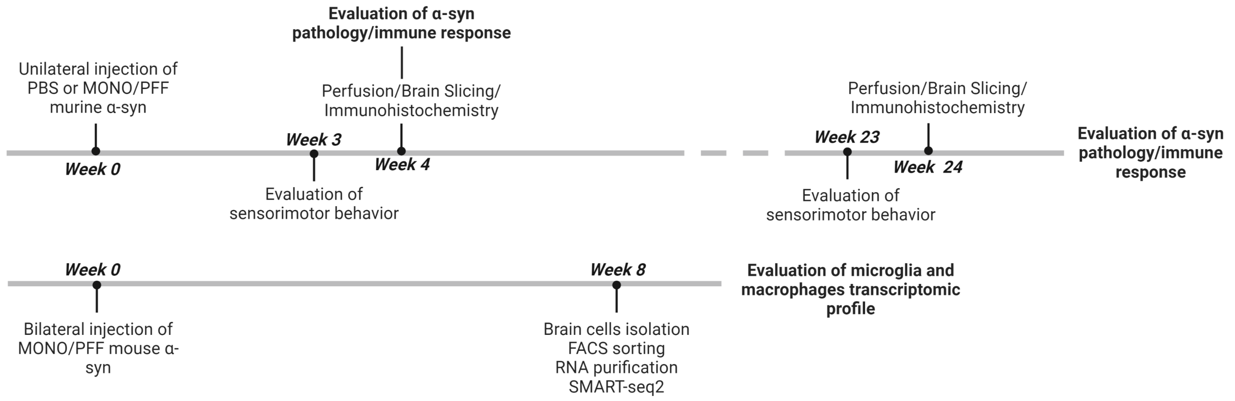


**Supplementary Fig. 1 Experimental study design.** Graphic Illustration of experimental study design for histology study and RNA sequencing study. Created with Biorender.com

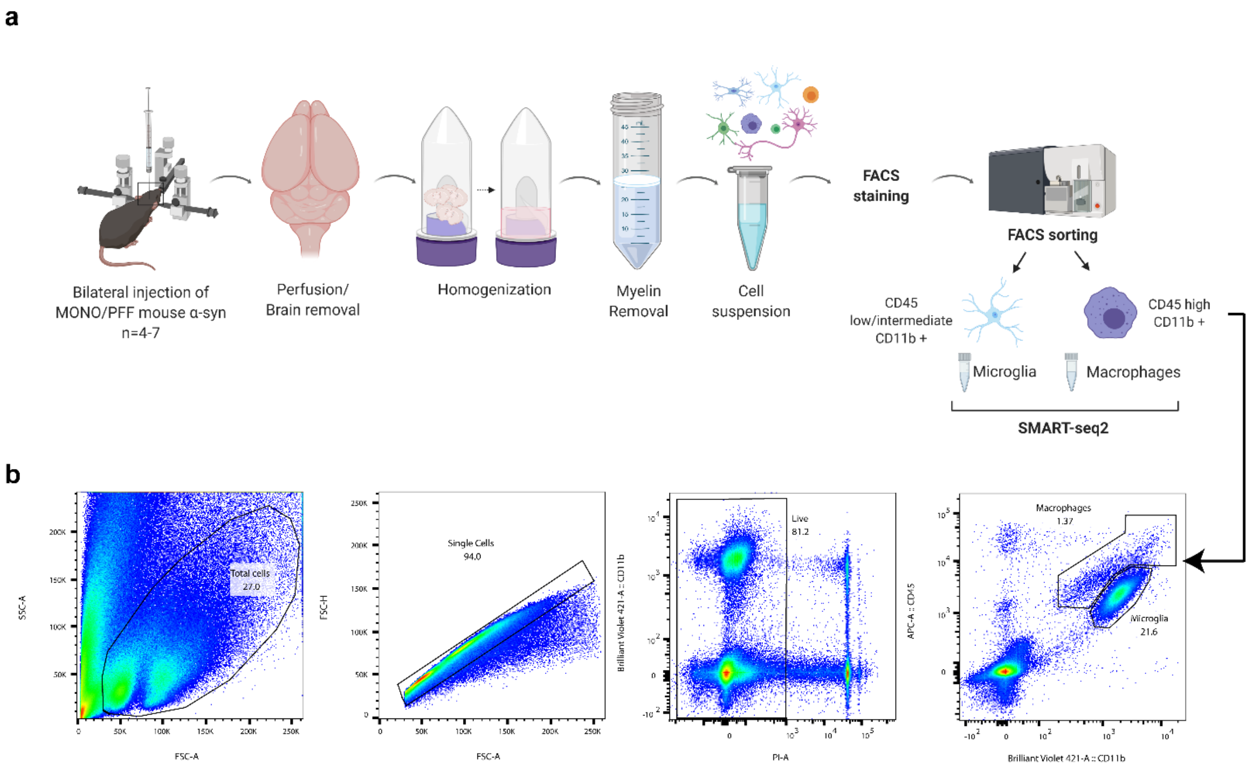


**Supplementary Fig. 2 Flow cytometry gating strategy for Microglia and Macrophage populations**. **a)** Image illustration of experimental study design of brain cells isolation for SMARTSeq2. **b)** Flow cytometry gating strategy for FACS sorting of Microglia (CD45 low/intermediate, CD11b positive) and Macrophage (CD45 high, CD11b positive) populations. Image created with BioRender.com

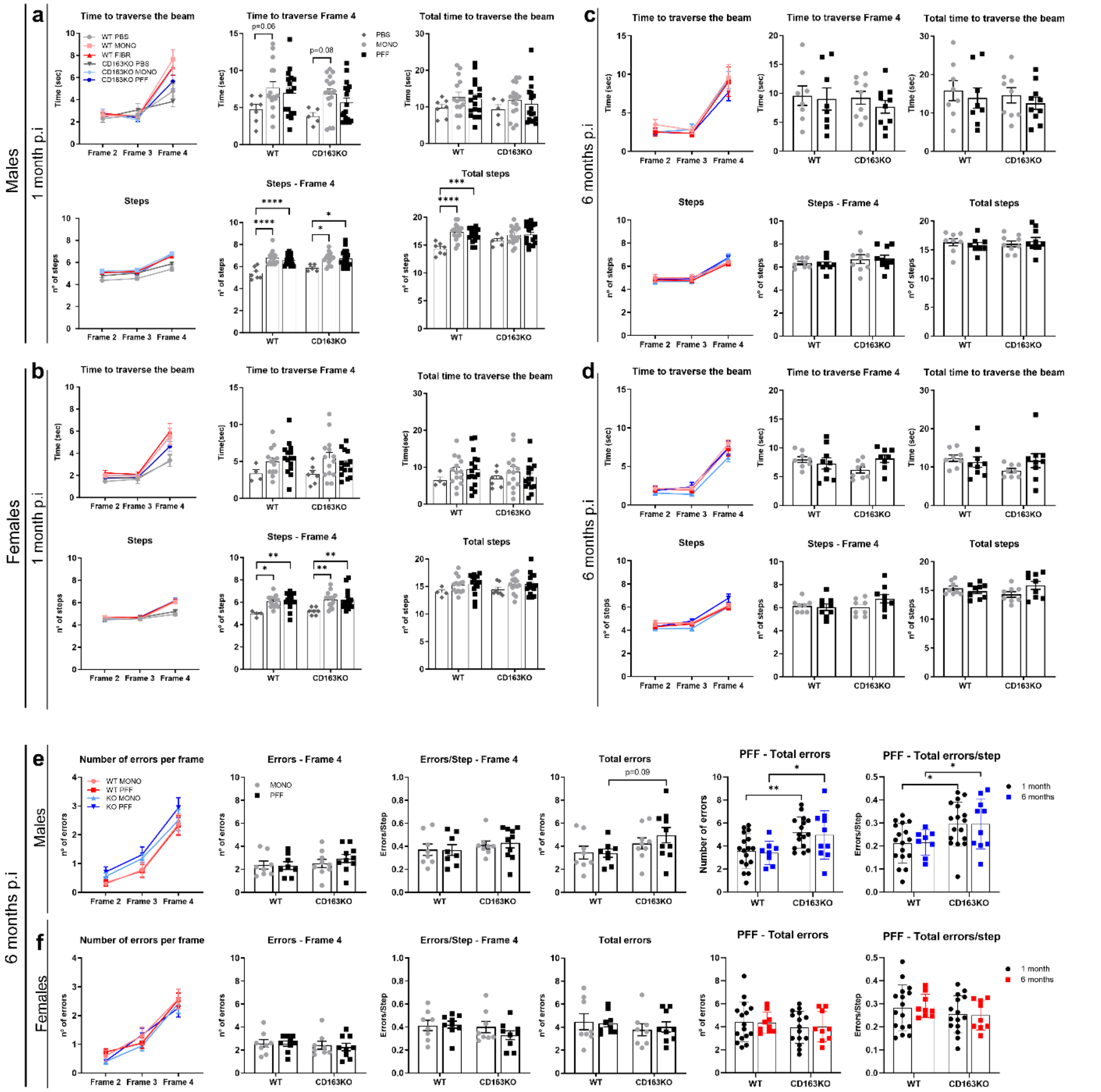


**Supplementary Fig. 3 Assessment of motor performance on the Challenging Beam test.** Line and bar graphs with individual values illustrate the quantitative measurement of the number of time and steps to traverse the beam, time and steps to traverse Frame 4 and total time/ total steps to traverse the beam (combination of Frame 2, 3 and 4) in the Challenging Beam test at **a-b)** 1-month post- PBS or α-syn MONO/PFF unilateral injection in the Striatum and **c-d)** 6-months post-α-syn MONO/PFF injection in **a, c)** males and **b, d)** females. Line and bar graphs with individual values illustrate the quantitative measurement of the number of errors per frame, number of errors and errors/step in Frame 4 and total number of errors (combination of Frame 2, 3 and 4) in the Challenging Beam test, 6-months post-α-syn MONO/PFF unilateral injection in the Striatum of **e)** males and **f)** females. Values are Mean±SEM (n=14-17 for behavior at 1-month; n=8-10 for behavior at 6-months). Statistics: Two-way ANOVA followed by Sidak's multiple comparison test.

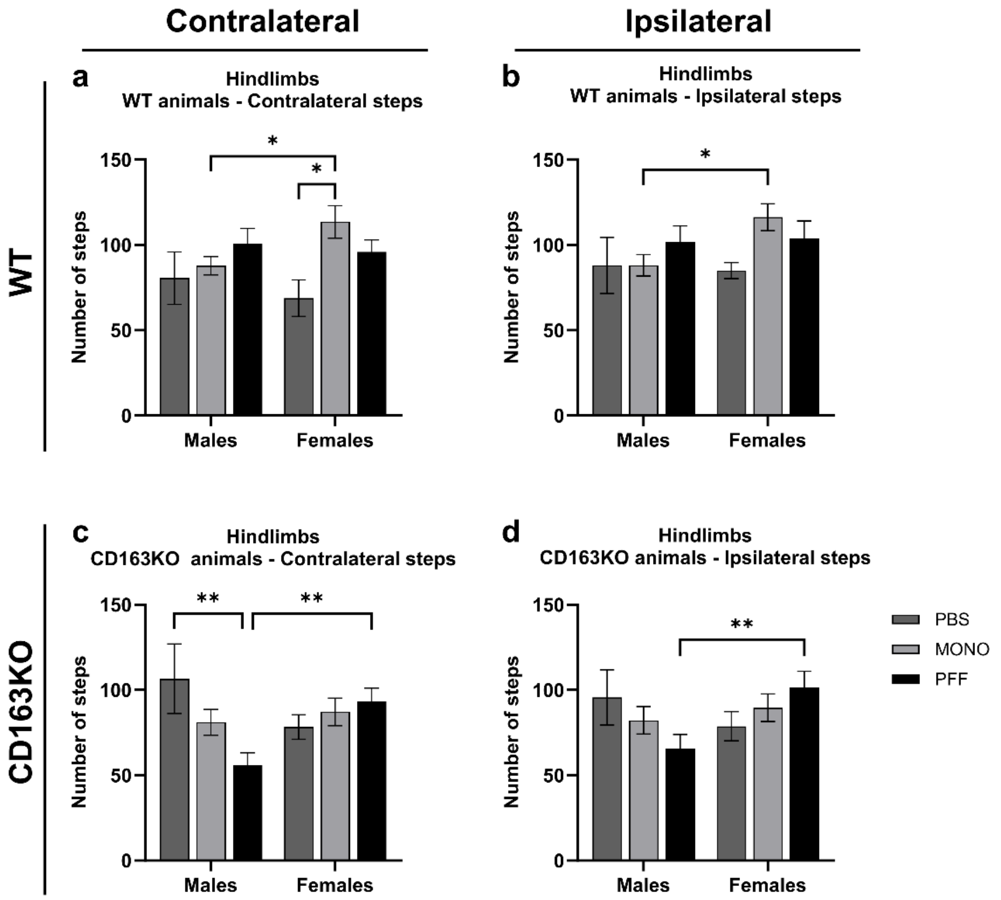


**Supplementary Fig.4 Assessment of motor performance on the Cylinder test 1 month post-injection.** Bar graphs represent the number of **a,c)** contralateral and **b,d)** ipsilateral hindlimb steps in the cylinder in **a-b)** WT and **c-d)** CD163KO males and females. Values are Mean±SEM (n=4-7 (PBS group), 14-17 (MONO/PFF groups) Statistics: Two-way ANOVA followed by Sidak's multiple comparison test. *p<0.05, **<0.01, ***<0.001, ****<0.0001.

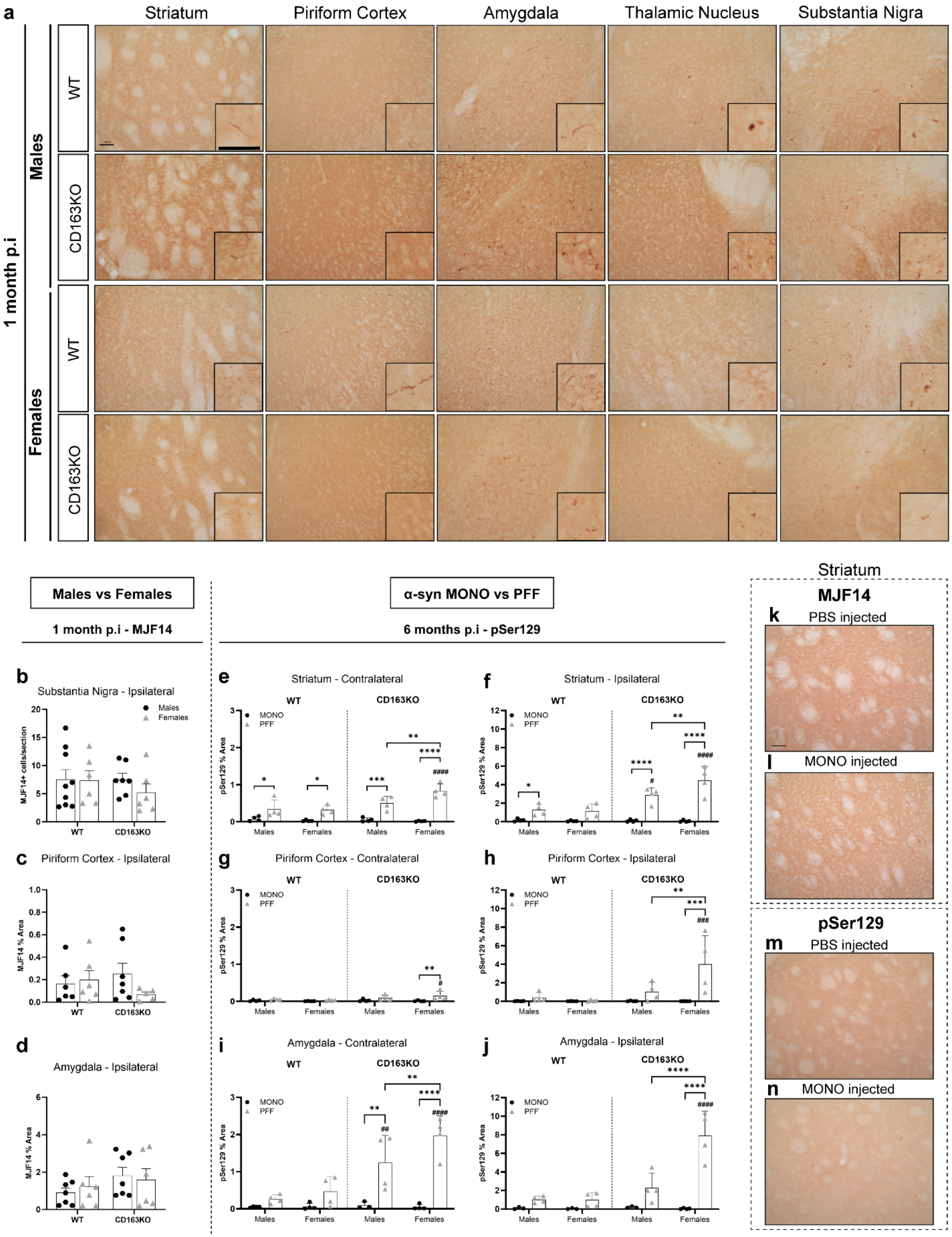


**Supplementary Fig.5 Spreading of pathological α-syn through the basal ganglia and interconnected regions. a)** Representative images of MJF14 immunostaining in α-syn PFF-injected animals at 1-month p.i. **b)** Bar graphs with average and individual points represent the number of MJF14+ cell aggregates in the ipsilateral substantia nigra, and the percentage of area covered by MJF14+ staining at 1-month post-α-syn-PFF injection in the **c)** ipsilateral piriform cortex and **d)** amygdala. **e-f)** Bar graphs with individual values represent the percentage of area covered by phosphorylated (pSer129) α-syn in the contralateral and ipsilateral striatum, **g-h)** piriform cortex and **i-j)** amygdala 6-month post-α-syn injection. Representative striatal images of **k-l)** MJF14 and **m-n)** pSer129 immunostaining in PBS and α-syn MONO-injected animals at 1-month p.i. Values are Mean±SEM (n=6-9 for MJF14) and (n=4 for pSer129). Statistics: Two-way ANOVA followed by Sidak's multiple comparison test. P values are given according to the number of symbols e.g. *p<0.05, **<0.01, ***<0.001, ****<0.0001. The symbol # indicates different to the corresponding WT group. Scale bar: 50µm applies to all.

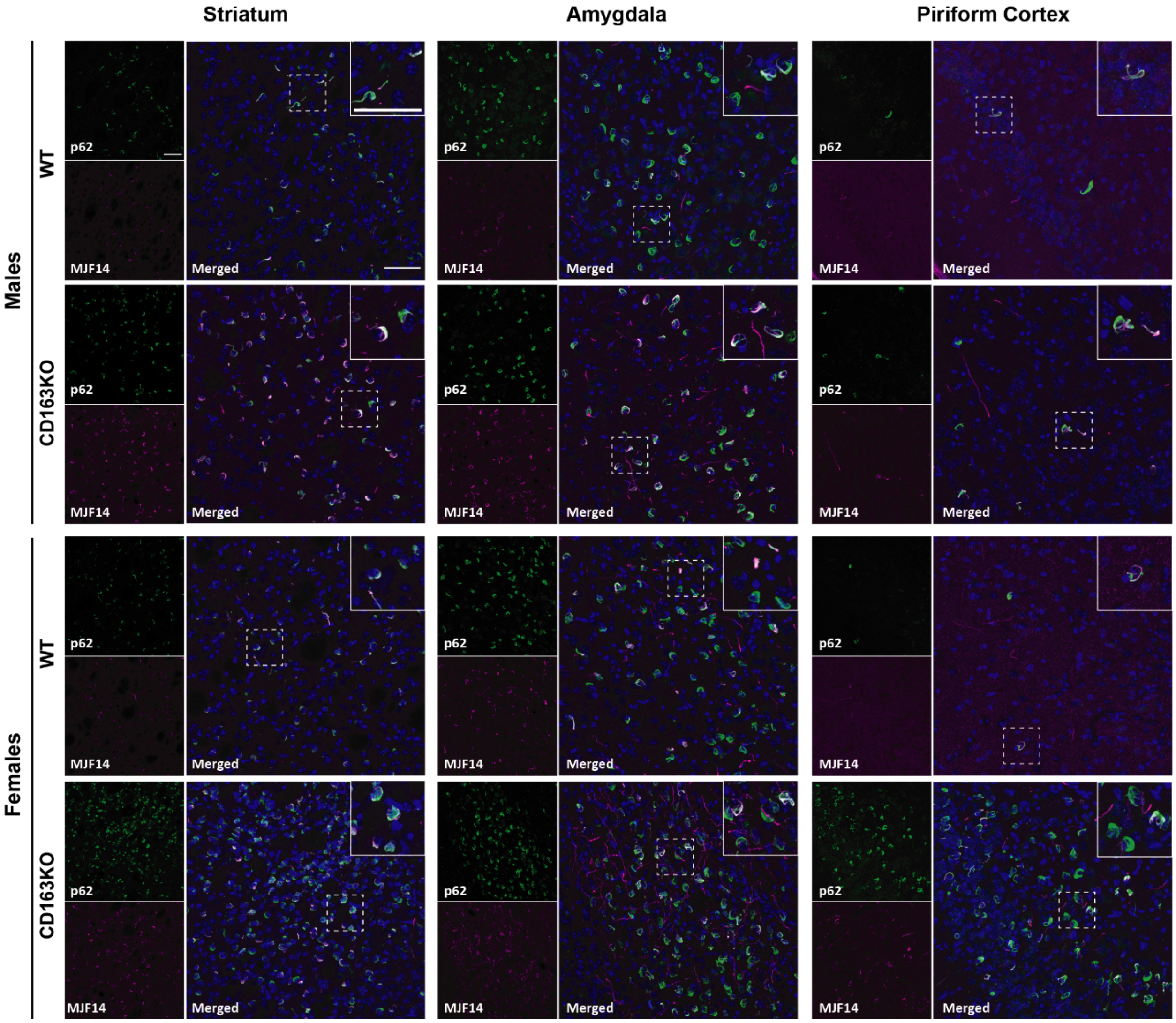


**Supplementary Fig.6 Aggregated α-syn is associated with p62 accumulation.** Representative confocal images of MJF14 (magenta) and p62 (green) and merged photo (with nuclear DAPI (blue)) in the ipsilateral striatum, amygdala and piriform cortex of α-syn PFF animals 6 months p.i. Squares with white dashed lines delineate the cropped region of the amplified images (upper right corner). Scale bar: 50 µm.

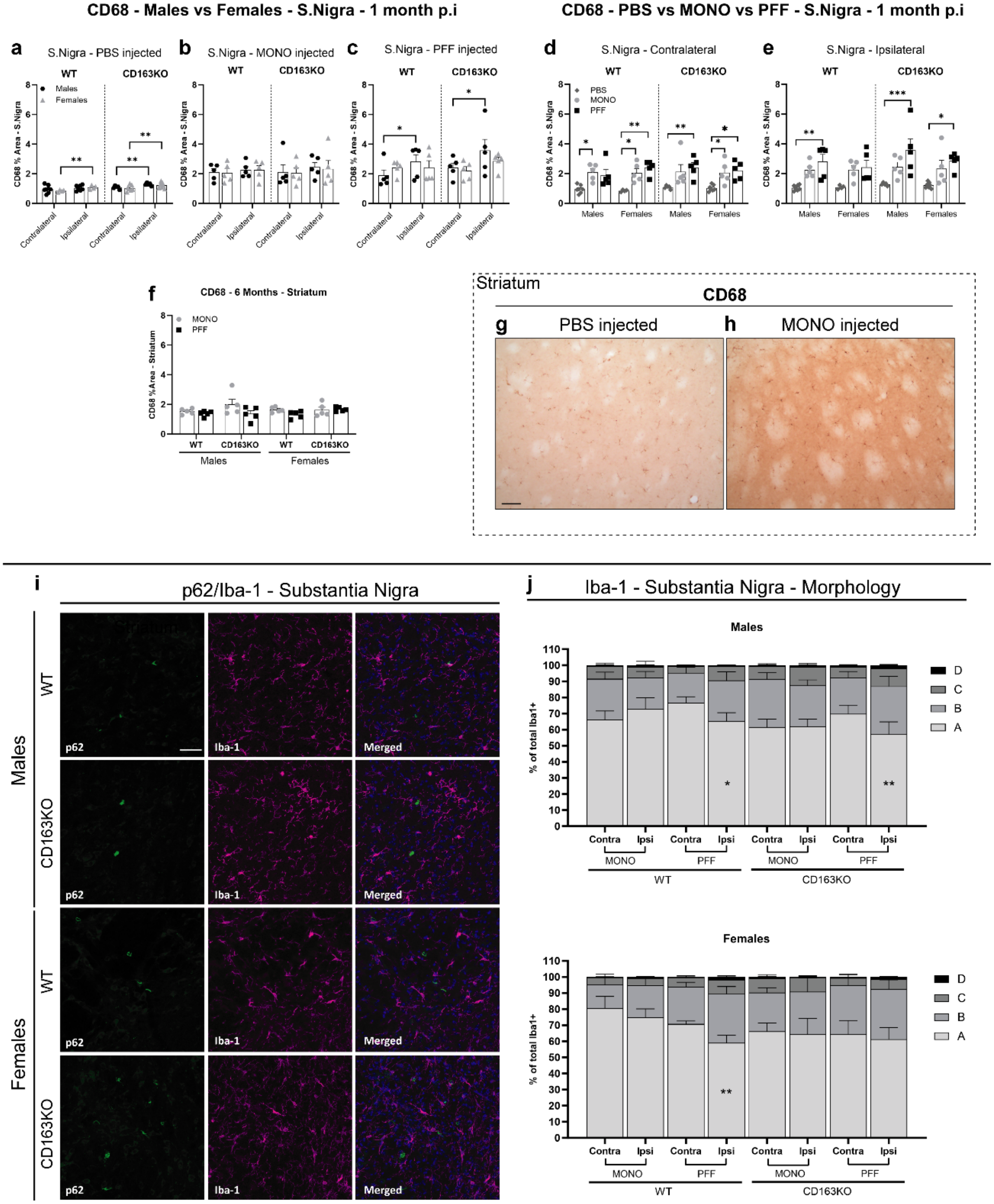


**Supplementary Fig.7 CD68 and Iba-1 expression after α-syn injection.** Bar graphs with individual values represent the percentage of area covered by CD68 staining in the **d)** contralateral and **e)** ipsilateral SN 1 month post- **a)** PBS **b)** α-syn MONO and **c)** α-syn PFF injection; and in the **f)** ipsilateral striatum 6 months post- α-syn MONO/PFF injection. Representative striatal images of CD68 immunostaining in **g)** PBS and **h)** α-syn MONO-injected animals at 1-month p.i. Representative confocal images of p62 (green) labeling with **i)** Iba-1 (magenta) at 6 months p.i, displaying individual channels and merged/composite images with (DAPI (blue)) in the ipsilateral SN of α-syn PFF animals. Scale bar: 50 µm applies to all. **j)** Stacked bar graphs illustrate the percentage of total Iba-1 positive cell subtypes (A,

B, C and D) in the contralateral and ipsilateral SN, 6 months after α-syn MONO and PFF-injection in Males (upper graph) and Females (lower graph). Values are Mean±SEM (n=6-9 for CD68) and (n=5 for Iba-1). Statistics: a-f) Two-way ANOVA followed by Sidak's multiple comparison test. *p<0.05, **<0.01, ***<0.001. **j)**: Two-way ANOVA (side and subtype) followed by Sidak's multiple comparison test. *p<0.05, **<0.01 (compared to contralateral in the same group and subtype).

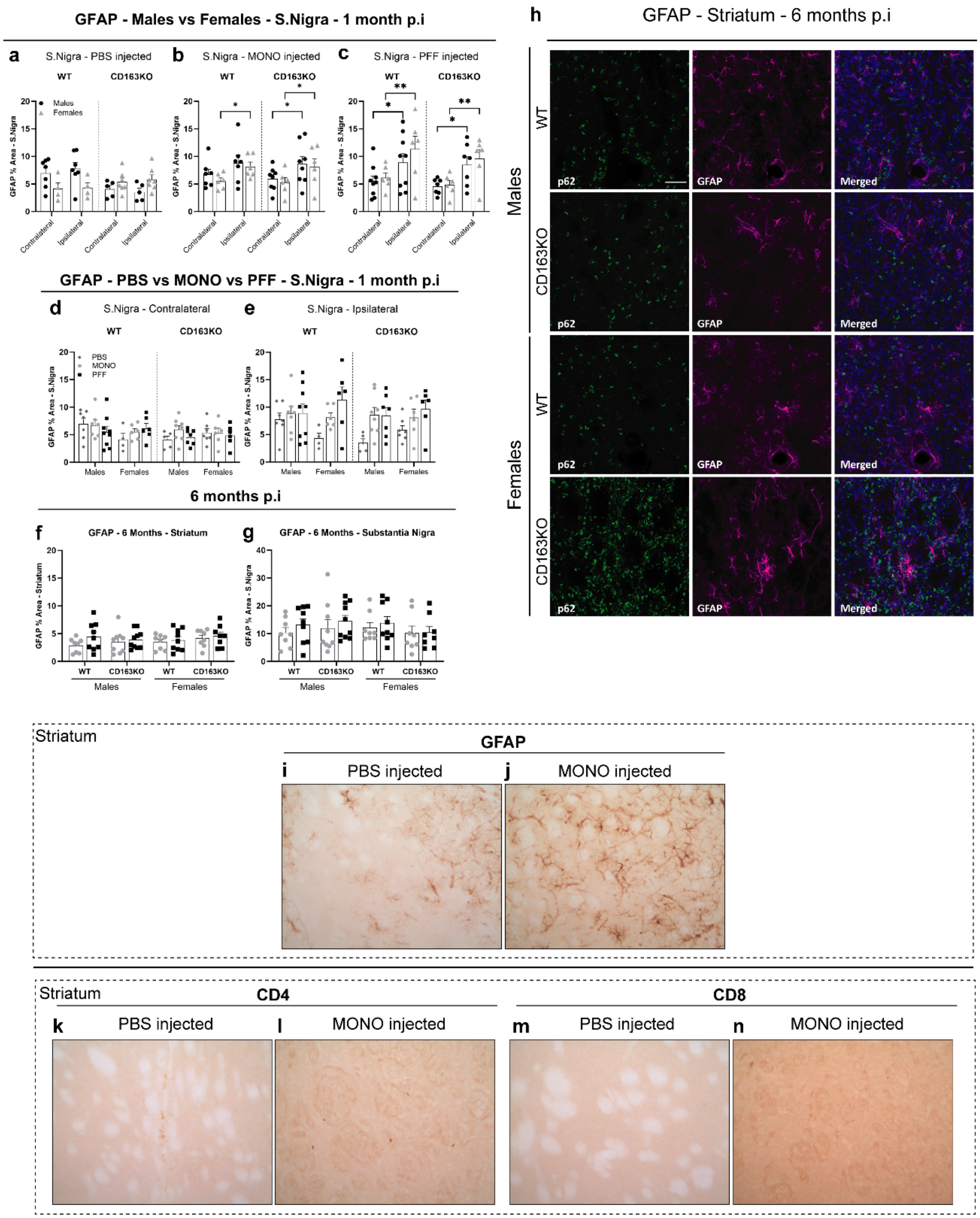


**Supplementary Fig.8 GFAP expression after α-syn injection.** Bar graphs with individual values represent the percentage of area covered by GFAP staining in the **d)** contralateral and **e)** ipsilateral SN 1 month post- **a)** PBS **b)** α-syn MONO and **c)** α-syn PFF injection; and in the **f)** ipsilateral striatum and **g)** ipsilateral SN 6 months post- α-syn MONO/PFF injection. Representative confocal images of p62 (green) labeling with **i)** GFAP (magenta) at 6 months p.i, displaying individual channels and merged/composite images (with DAPI (blue)) in the ipsilateral SN of α-syn PFF animals. Representative striatal images of GFAP, CD4 and CD8 immunostaining in **i,k,m)** PBS and **j,l,n)** α-syn MONO-injected animals at 1-month p.i.

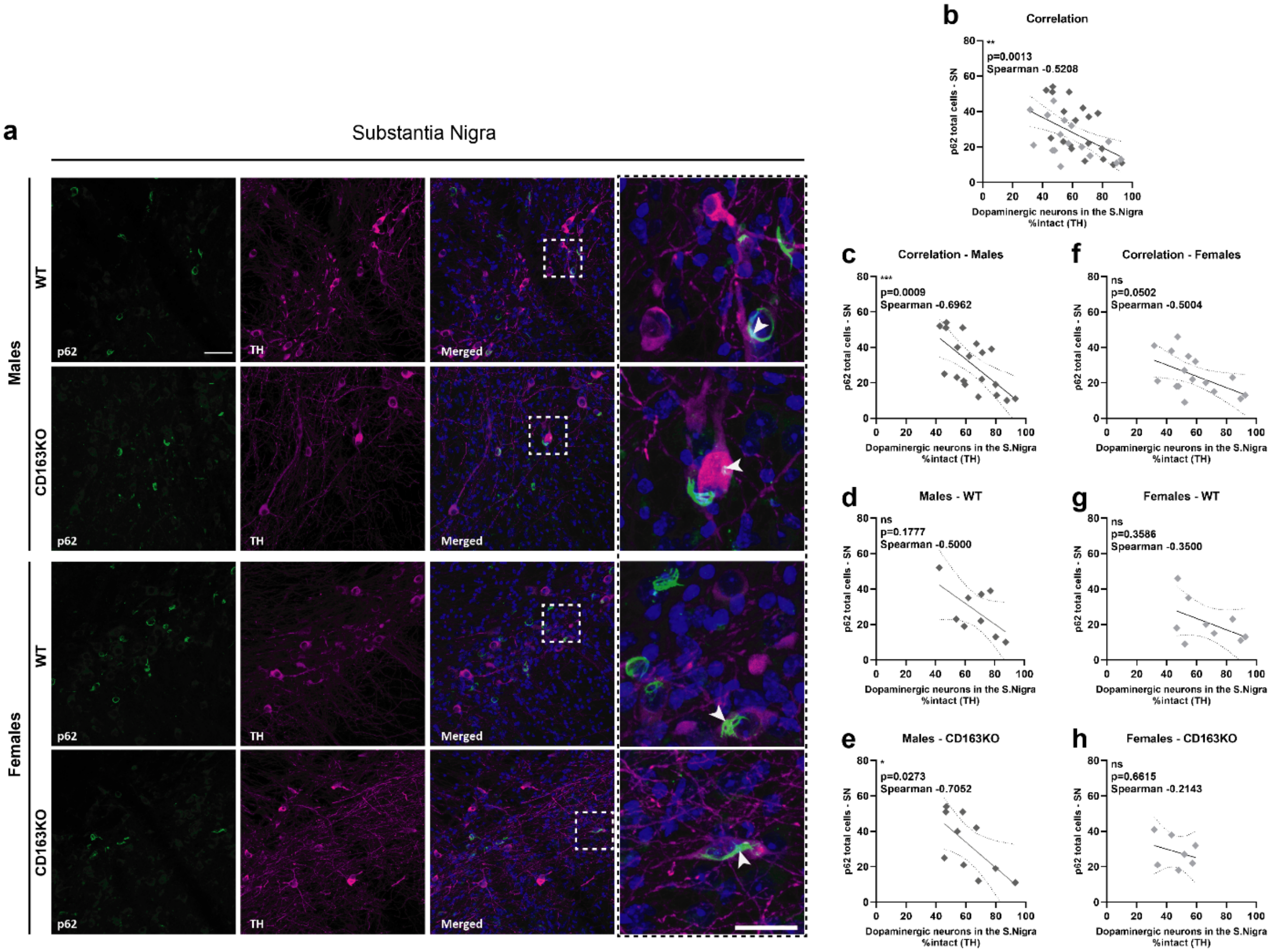


**Supplementary Fig.9 p62 positive inclusions co-localize with TH in SN cell bodies and are associated with cell death in CD163KO males. a)** Representative 20x view confocal images of TH (magenta) labeling with p62 (green) 6 months p.i, displaying individual channels and composite images (with DAPI (blue)) in the ipsilateral SN. Squares with white dashed lines delineate the cropped region of the amplified images. White arrowheads indicate co-localization. Scale bar represents 50 µm in the left panel images and 25 µm in the right cropped panel. **b-h)** Shows the correlation between the number of p62+ aggregates and the number of TH+ neurons in the SN of α-syn PFF animals. Statistics: Spearman two-tail p-values (*p<0.05, **<0.01, ***<0.001), Spearman r and best-fit slope with 95% confidence intervals are plotted.

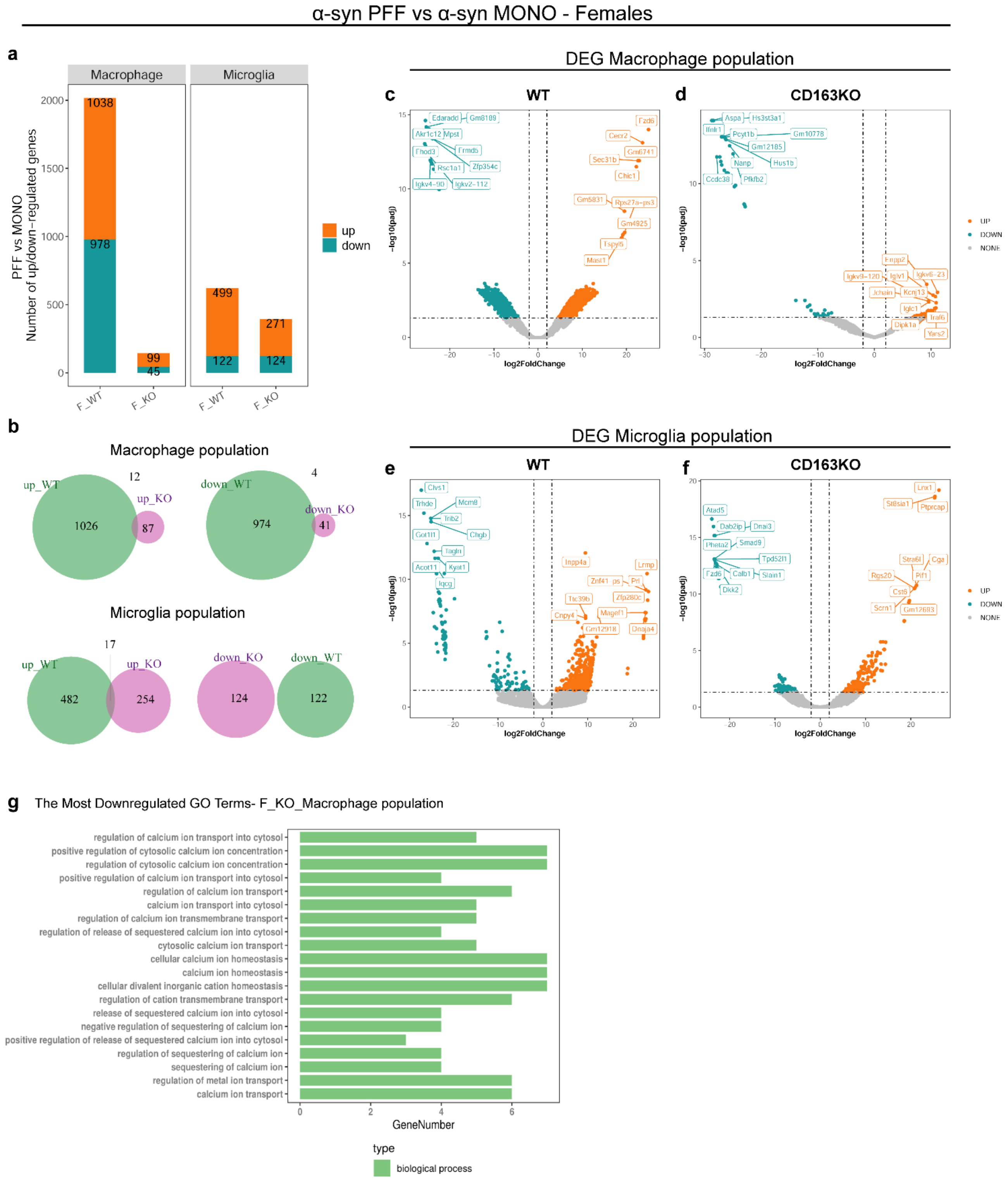


**Supplementary Fig.10. Differentially expressed genes (DEGs) in Female α-syn PFF Macrophage and Microglia populations (vs. MONO). a)** Bar graph showing the number of up/downregulated genes in α-syn PFF vs. MONO in Macrophage and Microglia populations 2 months p.i. **b)** Venn diagrams of shared DEGs. **c-f)** Volcano scatter-plots (-log10(padj) *vs* log2FoldChange) of DEGs in the Macrophage population and Microglia population. **g)** Gene Ontology (GO) analysis on the downregulated genes in CD163KO female macrophages.

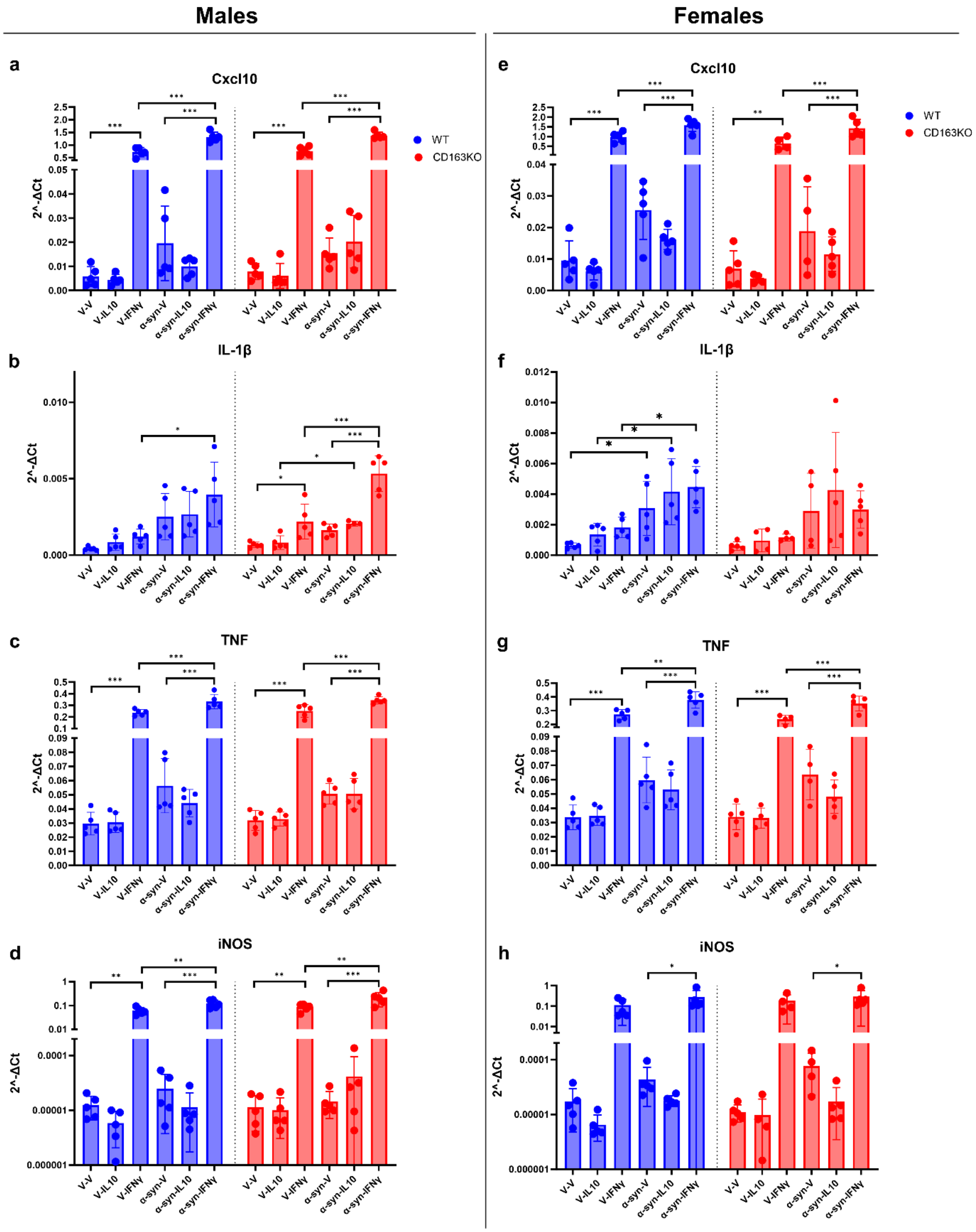


**Supplementary Fig.11 Pro-inflammatory gene expression profile of BMDM from WT and CD163KO mice after *in vitro* stimulation and α-syn PFF treatment.** Expression of M1 signature markers in *in vitro* stimulated BMDM isolated from WT and CD163KO **a-d)** males and **e-h)** females (n=4-5). BMDM were unstimulated (V=Vehicle) or stimulated with IL-10 and IFNγ 24h prior to α-syn PFF treatment (6h). Statistics: Values are Mean±SD. Two-way ANOVA or Mixed effects analysis with Bonferroni correction (<0.0071). *p<0.0071, **<0.00071, ***<0.000

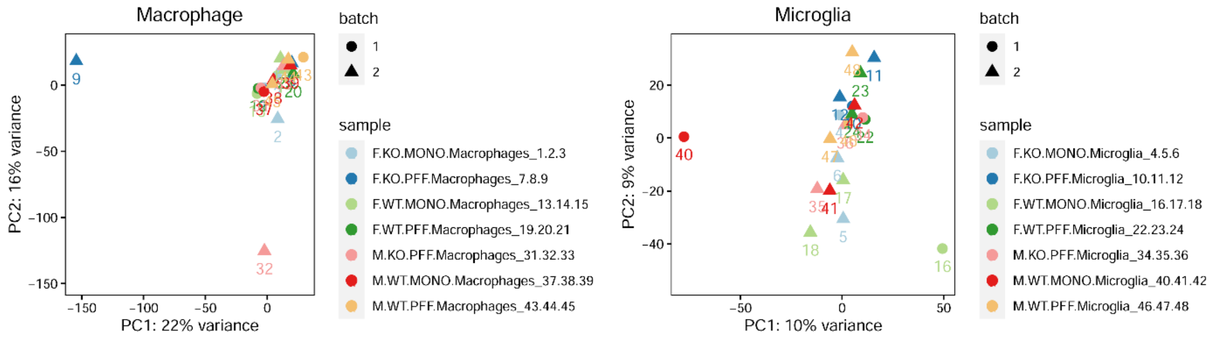


**Supplementary Fig. 12 Principal component analysis (PCA) on Macrophage and Microglia populations.** Batch effect was corrected and three outliers (Macrophage: #9 #32, Microglia: #16) were removed.
